# Imaging Sterols and Oxysterols in Mouse Brain Reveals Distinct Spatial Cholesterol Metabolism

**DOI:** 10.1101/450973

**Authors:** Eylan Yutuc, Roberto Angelini, Mark Baumert, Natalia Mast, Irina Pikuleva, Jillian Newton, Malcolm R. Clench, David O.F. Skibinski, Owain W. Howell, Yuqin Wang, William J. Griffiths

## Abstract

Dysregulated cholesterol metabolism is implicated in a number of neurological disorders. Many sterols, including cholesterol and its precursors and metabolites, are biologically active and important for proper brain function. However, spatial cholesterol metabolism in brain and the resulting sterol distributions are poorly defined. To better understand cholesterol metabolism *in situ* across the complex functional regions of brain, we have developed on-tissue enzyme-assisted derivatisation in combination with micro-liquid-extraction for surface analysis and liquid chromatography - mass spectrometry to image sterols in tissue slices (10 µm) of mouse brain. The method provides sterolomic analysis at 400 µm spot diameter with a limit of quantification of 0.01 ng/mm^2^. It overcomes the limitations of previous mass spectrometry imaging techniques in analysis of low abundance and difficult to ionise sterol molecules, allowing isomer differentiation and structure identification. Here we demonstrate the spatial distribution and quantification of multiple sterols involved in cholesterol metabolic pathways in wild type and *cholesterol 24S-hydroxylase* knock-out mouse brain. The technology described provides a powerful tool for future studies of spatial cholesterol metabolism in healthy and diseased tissues.

**Significance:** The brain is a remarkably complex organ and cholesterol homeostasis underpins brain function. It is known that cholesterol is not evenly distributed across different brain regions, however, the precise map of cholesterol metabolism in the brain remains unclear. If cholesterol metabolism is to be correlated with brain function it is essential to generate such a map. Here we describe an advanced mass spectrometry imaging platform to reveal spatial cholesterol metabolism *in situ* at 400 µm resolution on 10 µm tissue slices from mouse brain. We mapped, not only cholesterol, but also other biologically active sterols arising from cholesterol turnover in both wild type and mice lacking cholesterol 24-hydroxylase (Cyp46a1), the major cholesterol metabolising enzyme.

## 1. Introduction

The brain is a remarkably complex organ in terms of anatomy and function. Little is known about the landscape of the metabolome or lipidome across the brain. The brain represents a major repository of unesterified cholesterol in mammals, almost 25% of total body cholesterol is found in brain and the central nervous system (CNS), where it is present at a level of about 20 µg/mg (wet weight) (1). As cholesterol cannot cross the blood brain barrier (BBB) and be imported from the periphery, essentially all cholesterol in brain is biosynthesised in brain. Inborn errors of cholesterol biosynthesis often result in neurodevelopmental disorders and retardation syndromes (2). It is recognized that not only the steady-state level of cholesterol is critical for brain function but also its constant turnover (3, 4). Much of brain cholesterol is found in myelin sheaths of oligodendrocytes surrounding the axons of neurons, where it is turned over slowly (0.4% per day in mouse, 0.03% in human (1)), but a more metabolically active pool of cholesterol is located in intracellular structures such as the endoplasmic reticulum, Golgi apparatus and nucleus of neurons. Here turn-over is estimated at 20% per day in mouse (1). However, a precise map of cholesterol turnover across the specialized functional regions of brain remains to be generated.

Oxysterols are oxidised forms of cholesterol, or of its precursors, and represent the initial metabolites in cholesterol catabolism (5). One of the most well studied oxysterols in brain is 24S-hydroxycholesterol (24S-HC, See SI Appendix, Table S1 for abbreviations, common and systematic names). In mammals 24S-HC is mostly synthesised in neurons by the enzyme cytochrome P450 (CYP) 46A1 (cholesterol 24S-hydroxylase, CYP46A1) and acts as a transport form of cholesterol (SI Appendix, Figure S1) (4, 6, 7). Unlike cholesterol, 24S-HC can cross the BBB. 24S-HC is a ligand to the liver X receptors (LXRs) (8), the β-form of which is highly expressed in brain (9), a potent modulator of the *N*-methyl-D-aspartate receptors (NMDARs) (10), glutamate-gated ion channels that are critical to the regulation of excitatory synaptic function in the CNS, and is also a modulator of cholesterol biosynthesis by interacting with the endoplasmic reticulum protein INSIG (insulin induced gene) and restricting transport and processing of SREBP-2 (sterol regulatory element-binding protein-2) to its active form as a master transcription factor for cholesterol biosynthesis (11). Despite its potent biological activity, little is known about levels of 24S-HC in distinct regions of brain (12, 13). This is also true of 24S,25-epoxycholesterol (24S,25-EC), which is formed via a shunt pathway of cholesterol biosynthesis in parallel to the Bloch pathway but without the involvement of 24-dehydrocholesterol reductase (DHCR24, SI Appendix, Figure S1) (14, 15). 24S,25-EC has also been shown to be formed from desmosterol by CYP46A1 *in vitro* (16). 24S,25-EC is a potent activator of the LXRs, inhibitor of SREBP-2 processing and also a ligand to the G protein-coupled receptor (GPCR) Smoothened (SMO), a key protein in the Hedgehog (Hh) signalling pathway (8, 11, 17, 18). 24S,25-EC has been shown to be important in dopaminergic neurogenesis via activation of LXRs (19, 20).

The cholestenoic acids, 3β,7α-dihydroxycholest-5-en-(25R)26-oic (3β,7α-diHCA) and 7α-hydroxy-3-oxocholest-4-en-(25R)26-oic (7αH,3O-CA) acids, also formed by enzymatic oxidation of cholesterol, are found in cerebrospinal fluid (CSF), the fluid that bathes the CNS (21, 22). 7αH,3O-CA has been suggested to provide an export route for (25R)26-hydroxycholesterol (26-HC, also called 27-hydroxycholesterol) which itself passes into the brain from the circulation, but does not accumulate in brain (see SI Appendix, Figure S1) (23, 24). 3β,7α-diHCA has been shown to be biologically active as a ligand to the LXRs and to be protective towards oculomotor neurons (21). The location of the acid has not previously been defined in brain (25).

To understand better the importance in brain of sterols in general, and oxysterols in particular, it is necessary to correlate molecular concentrations with histology. This can be achieved by exploiting mass spectrometry imaging (MSI). MSI technology mostly utilises matrix-assisted laser desorption/ionisation (MALDI)-MS (26–29). MALDI-MSI has been used to image lipids in brain (30–32), however, cholesterol and other sterols tend to be poorly ionised by conventional MALDI and are underrepresented in MALDI-MSI studies. To enhance ionisation other desorption/ionisation methods have been employed, including nanostructure-initiator MS (33), sputtered silver-MALDI (34) and silver nanoparticle-MALDI (35, 36). These studies are mainly restricted to cholesterol. Alternatively, on-tissue derivitisation has been used to enhance MS-ionisation and to image steroid hormones and steroidal drugs in tissue by exploiting hydrazine reactions with carbonyl groups present in target analytes (e.g. Figure 1A) (37–42). These derivatives have been used in combination with MALDI and with “liquid extraction for surface analysis” (LESA) technology, or with reactive-DESI (desorption electrospray ionisation) (37, 43). Hydrazine derivatisation is problematic for sterols, oxysterols or steroids not possessing a carbonyl group. To address this issue, for in-solution analysis of oxysterols and sterols, we and others have utilised a bacterial cholesterol oxidase enzyme to specifically convert sterols with a 3β-hydroxy-5-ene structure to a 3-oxo-4-ene and then derivatise the resulting 3-oxo group with the Girard hydrazine reagent to provide a charge-tag to the target molecule (Figure 1C) (44–46). This strategy is termed enzyme-assisted derivatisation for sterol analysis (EADSA). For comprehensive analysis, oxysterols naturally possessing an oxo group, e.g. 7αH,3O-CA or 7-oxocholesterol (7-OC, also called 7-ketocholesterol) can be derivatised in parallel in the absence of cholesterol oxidase (Figure 1B).

**Figure 1.**
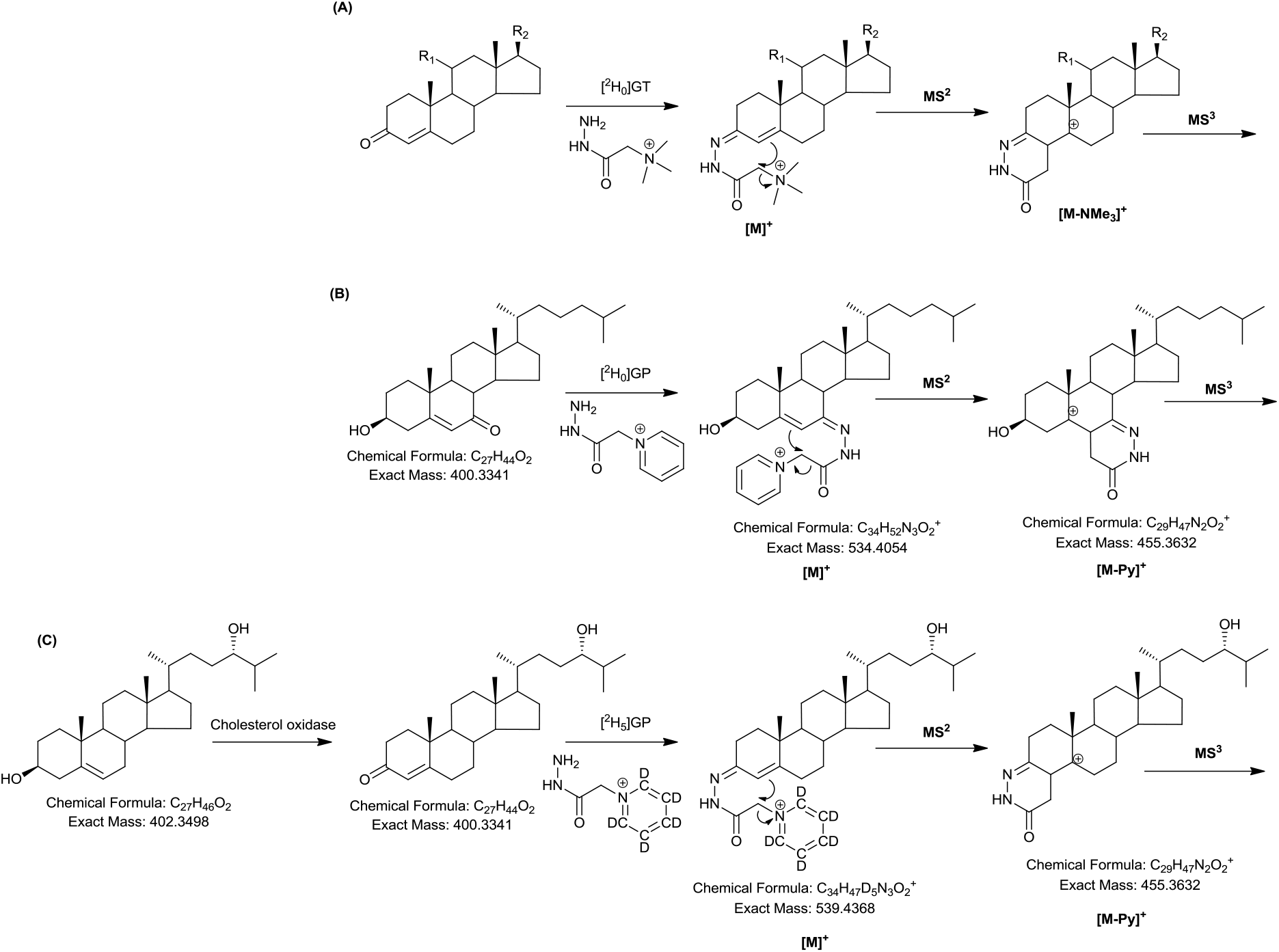
Derivatisation and fragmentation of steroids, sterols and oxysterols. (A) Derivatisation of corticosteroids (corticosterone, R_1_ = OH, R_2_ = COCH_2_OH; 11-dehydrocorticosterone, R_1_ = O, R_2_ = COCH_2_OH) or testosterone (R_1_ = H, R_2_ = OH) with Girard T (GT) reagent. (B) Derivatisation of 7-OC with [^2^H_0_]Girard P reagent ([^2^H_0_]GP) and of (C) 24S-HC with [^2^H_5_]GP reagent following enzymatic oxidation with cholesterol oxidase.

In the current study we describe how on-tissue EADSA with micro-scale-LESA (µLESA) and liquid chromatography (LC) - MS can be exploited for the imaging of cholesterol, its precursors and metabolites, in rodent brain. The incorporation of on-line LC is essential to separate isomers and also low abundance sterols from dominating cholesterol. We find 24S-HC to be the most abundant oxysterol in all brain regions, being at highest levels in striatum and thalamus and at lowest levels in grey matter of cerebellum. 24S,25-EC is also abundant in thalamus and sparse in grey matter of cerebellum. We are able to definitively identify 3β,7α-diHCA in brain, where it is most abundant in grey matter of the cerebellum. We confirm that 24S-HC is almost completely absent from brain of the *Cyp46a1* knockout mouse (*Cyp46a1*-/-). The level of 24S,25-EC is reduced in the *Cyp46a1*−/− mouse but low levels of 20S-hydroxycholesterol (20S-HC), 22R-hydroxycholesterol (22R-HC), 24R-hydroxycholesterol (24R-HC) and 25-hydroxycholesterol (25-HC) are present in this mouse.

## 2. Results

### 2.1. Development of Analytical Platform

The platform was designed to allow the MSI of cholesterol, its immediate precursors, and of multiple oxysterols in tissue against a large background of cholesterol and other more readily ionised lipids. To meet this challenge, we have used on-tissue derivatisation (Figure 1C) in combination with µLESA and LC-MS with multistage fragmentation (MS^n^). Each of the system components required optimisation as described below.

#### 2.1.1. On-Tissue EADSA

The protocol for on-tissue enzymatic oxidation of sterols (Figure 1C) was adapted from the in-solution procedure using cholesterol oxidase enzyme in KH_2_PO_4_ buffer (44, 46), the only difference being that the enzyme was sprayed onto tissue in buffer and then incubated at 37 °C in a humid chamber (11.9 cm x 8.2 cm x 8.5 cm, 30 mL water), rather than being in solution. In trial experiments, after a 1 hr incubation period at 37 °C there was no evidence for any non-oxidised sterol, as indicated by an absence of peaks corresponding to [M+H]^+^, [M+Na]^+^, [M+K]^+^, [M+H-(H_2_O)_n_]^+^, [M+Na-(H_2_O)_n_]^+^ or [M+K-(H_2_O)_n_]^+^ of endogenous cholesterol or 24S-HC, or of added deuterated-standards, only of the 3-oxo-4-ene oxidation products (see also SI Appendix). The next step in the method was on-tissue derivatisation of oxo-containing sterols using the Girard P (GP) hydrazine reagent (Figure 1C). Others have used Girard T (GT) hydrazine as the derivatising agent to react on-tissue with oxo-containing sterols/steroids (37–42), however, based on our experience of using the GP reagent for in-solution derivatisation of oxysterols, GP hydrazine was preferred here (44, 46). A disadvantage of using GT hydrazine is that a major MS^2^ fragment-ion is the [M-59]^+^ species (Figure 1A, [M-NMe_3_]^+^) and the exact same neutral-loss is observed in the fragmentation of endogenous choline-containing lipids. With [^2^H_0_]GP the major MS^2^ fragment-ion is [M-79]^+^ and for [^2^H_5_]GP [M-84]^+^, neutral-losses not common for other lipids. We sprayed GP reagent (6.3 mg/mL for [^2^H_5_GP bromide) in the same solvent as previously used for in-solution derivatisation (70% methanol, 5% acetic acid) (44, 46). As in reported studies using the GT reagent (37, 38, 41), it was essential to incubate the GP-coated tissue in a humid atmosphere to achieve efficient derivatisation. Initial tests in a dry atmosphere revealed no derivatisation of 3-oxo-4-ene substrates. Incubation in a humid chamber (12 cm x 12 cm x 7.2 cm) containing 30 mL of 50% methanol, 5% acetic acid at 37°C for 1 hr provided an efficient derivatisation environment with minimum lateral dispersion of analytes (see SI Appendix, Figure S2). Increasing the volume or organic content of solution led to lipid delocalisation, while reducing the volume and organic content provided less efficient derivatisation based on ion-current in µLESA-MS measurements.

#### 2.1.2. µLESA

LESA is based on liquid micro-junction surface sampling (47), where solvent is deposited as a small volume liquid micro-junction by a robotic sampling probe, i.e. pipette tip or capillary, onto a surface, aspirated back to the tip or capillary and the extract analysed by electrospray ionisation (ESI)-MS. The LESA^PLUS^ configuration of the TriVersa Nanomate incorporates a 200 µm i.d. / 360 µm o.d. fused silica capillary replacing the conventional pipette tip (48) reducing the extraction spot size to ≤1 mm (49, 50). However, when using a 50% methanol solvent, as required here for extraction of derivatised sterols, we were unable to achieve this spot size as the surface tension required for a stable liquid micro-junction was not attainable. To overcome this problem, the sampling probe can be used to make a seal against the tissue surface, thereby minimising the possibility of solvent spreading (51). We have attached a 330 µm i.d. / 794 µm o.d. FEP (fluorinated ethylene propylene) sleeve to the fused silica capillary and used it to make a direct seal with the tissue surface preventing spreading of the extraction solvent beyond the boundary of the sleeve (see SI Appendix, Figure S3A). This gave a spot size of diameter ≤400 µm, as assessed by microscopic evaluation of tissue after liquid extraction performed in triplicate (see SI Appendix, Figure S3B). A 50% methanol extraction solvent was chosen to closely match the derivatisation solvent. The volume of solvent dispensed onto (more correctly, forced to be in contact with) tissue, was optimised to maximise oxysterol extraction with an extraction time of 30 s, as assessed by MS ion-current for target oxysterols (e.g. 24S-HC), without compromising spatial resolution i.e. leaking of solvent through the tissue – FEP seal. The number of repeat extractions on the same spot was similarly optimised. The optimal conditions were a “dispensed volume” of 1 µL, an extraction time of 30 s, performed three times on the same spot.

#### 2.1.3. EADSA-µLESA-LC-MS

Previous studies using MS have defined concentrations of 24S-HC, the major oxysterol in mouse brain, to be of the order of 20 – 60 ng/mg (wet weight) (25, 52–54). For comparison, the concentration of cholesterol is in the range of 10 – 20 µg/mg (1, 25, 53), and after EADSA treatment, in the absence of chromatographic separation, cholesterol totally dominates the µLESA-MS analysis of the mouse brain surface extract (Figure 2A). However, with the addition of LC to give the µLESA-LC-MS platform, analysis of 400 µm diameter spot extracts of brain tissue give clear separation of peaks corresponding to 24S-HC, 24S,25-EC, desmosterol, 8(9)-dehydrocholesterol (8-DHC) and cholesterol (Figure 2B). 8-DHC is an enzymatically generated isomer of 7-dehydrocholesterol (7-DHC), the immediate precursor of cholesterol in the Kandustch-Russell pathway (55).

**Figure 2.**
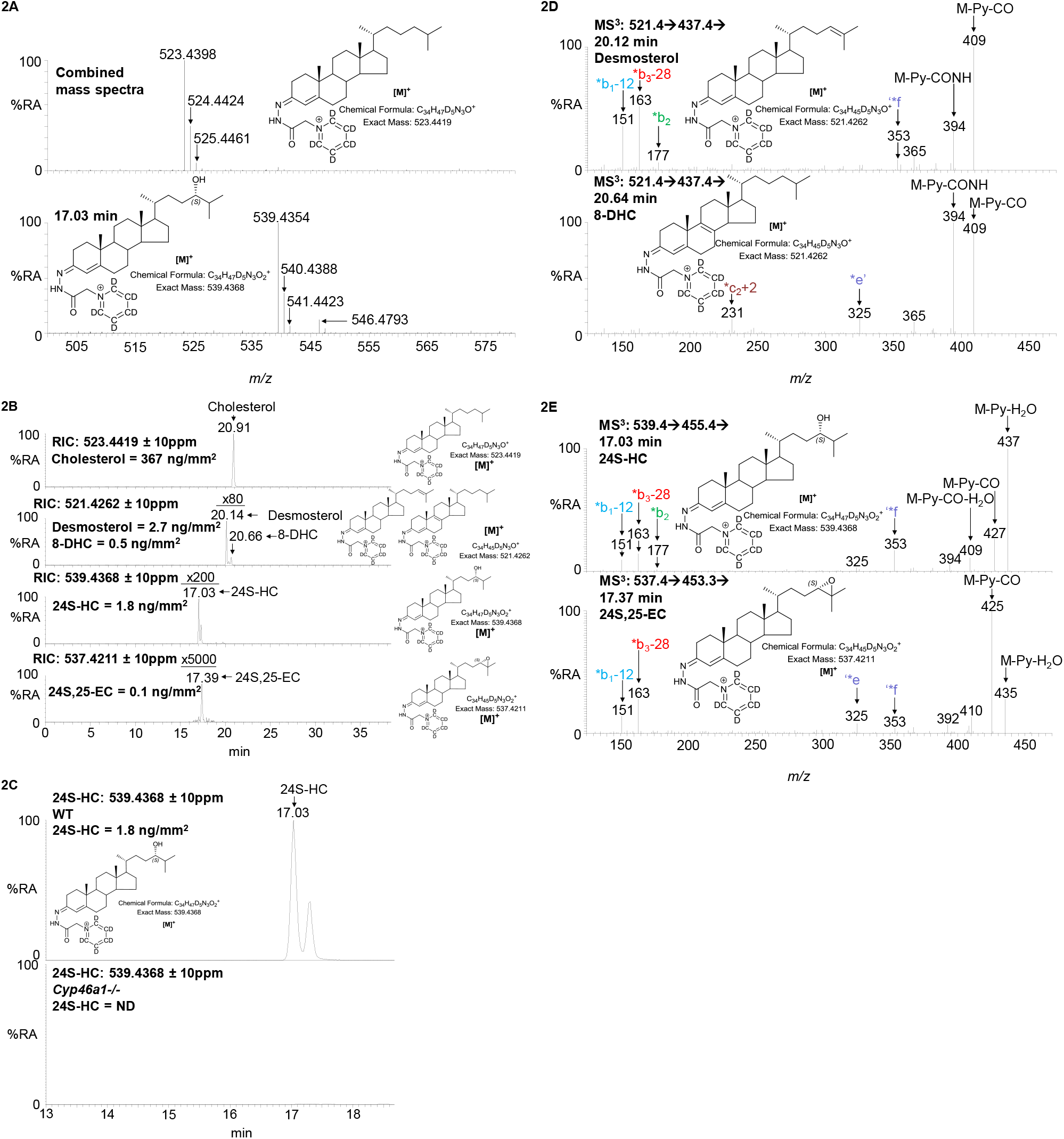
µLESA-LC-MS(MS^n^) analysis following EADSA treatment of mouse brain. (A) Upper panel, combined mass spectra recorded over the entire chromatogram, from striatum of WT mouse brain. Lower panel, mass spectrum recorded at 17.03 min from the same brain region. The peaks at *m/z* 523.4398, 539.4354 and 546.4793 correspond to endogenous cholesterol, 24S-HC and [^2^H_7_]24S-HC internal standard, respectively. (B) RICs for the [M]^+^ ion of endogenous cholesterol (*m/z* 523.4419 ± 10 ppm, top panel), dehydrocholesterols (521.4262 ± 10 ppm, 2^nd^ panel), 24S-HC (539.4368 ± 10 ppm, 3^rd^ panel) and 24S,25-EC (537.4211 ± 10 ppm, bottom panel) from striatum of WT mouse brain. Peak intensities are relative to cholesterol but magnified as indicated. (C) RIC for the [M]^+^ ion of endogenous 24S-HC (*m/z* 539.4368 ± 10 ppm) from striatum of WT brain (upper panel) and of *Cyp46a1−/−* mouse brain (lower panel). Peak intensities are normalised to the 24S-HC peak from WT mouse brain. ND, not detected. Note, 24S-HC appears as a doublet in chromatograms in (B) and (C), this is a consequence of *syn* and *anti* conformers of the GP-derivative. (D) MS^3^ ([M]^+^→[M-Py]^+^→) spectra recorded at 20.12 min corresponding to endogenous desmosterol (upper panel) and 20.64 min corresponding to 8-DHC (lower panel) from striatum of WT mouse. (E) MS^3^ ([M]^+^→[M-Py]^+^→) spectra recorded at 17.03 min corresponding to endogenous 24S-HC (upper panel) and 17.37 min corresponding to 24S,25-EC (lower panel) from striatum of WT mouse. Note in the EADSA process 24S,25-EC isomerises to 24-oxocholesterol.

We integrated µLESA with two different LC-MS systems; (i) µLESA-LC-MS with a “conventional” flow rate of 200 µL/min, and (ii) µLESA-nano-LC-MS with a flow rate of 350 nL/min. Both systems incorporate a micro-switch and a reversed-phase trap column prior to an analytical column which allowed trapping of GP-derivatised sterols and washing to waste of unreacted GP reagent and more polar analytes (Figure 3 and SI Appendix, Figure S4). The nano-LC-MS format gives a 10-fold improvement in sensitivity over the “conventional” flow-rate LC-MS format, based on analysis of 24S-HC, and is more suitable for the analysis of very low abundance oxysterols, however, it is susceptible to overloading by highly abundant cholesterol. Both systems were utilised in this study to cover the broadest range of sterols in brain.

**Figure 3.**
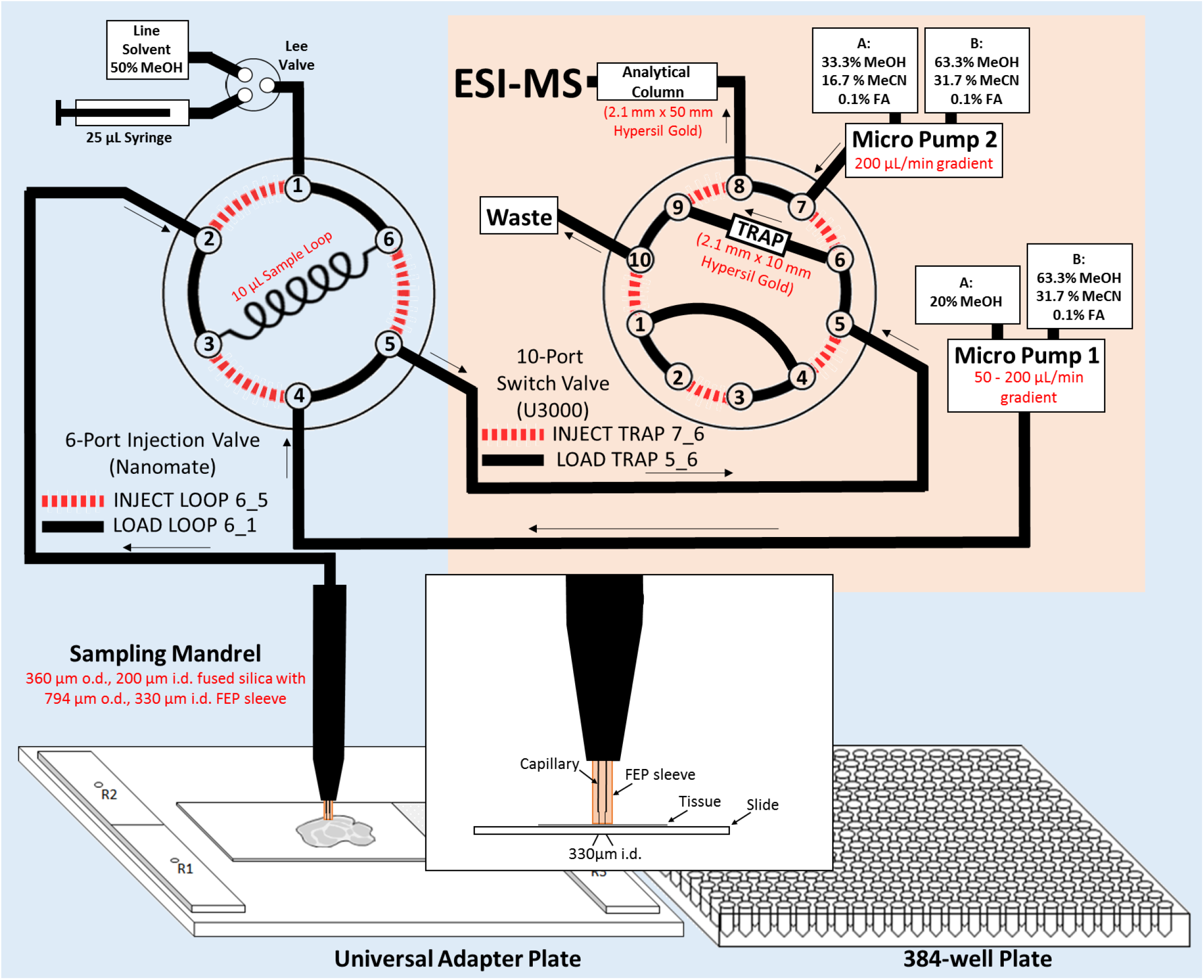
Schematic representation of the of the µLESA system linked to with LC-MS operating at conventional flow-rates. With connections between ports 1 and 6 and between 3 and 2, the 6-port injection valve is in the load position. Once loaded, connections are made between ports 4 and 3, and between 6 and 5. Sample is transported to the 10-port valve and with connections made between ports 5 and 6 and 9 and 10 (on the 10-port valve), sterols are trapped on the C_18_ trap-column. Port 7 is then connected to 6, port 9 to 8 and sterols eluted from the trap column to the analytical LC column. A schematic illustrating the set-up for nano-LC-MS is shown in SI Appendix, Figure S4.

#### 2.1.4. Quantification Using On-Tissue Internal Standards

To quantify sterols and oxysterols isotope-labelled surrogates were sprayed on-tissue providing known areal densities of internal standards. Initially, the validity of using sprayed-on [^2^H_7_]24R/S-HC as an internal standard for quantification of oxysterols was assessed by spraying on both [^2^H_7_]24R/S-HC and [^2^H_7_]22S-HC onto six successive slices of brain tissue at increasing concentration ratios of [^2^H_7_]24R/S-HC: [^2^H_7_]22S-HC, but keeping the areal density of [^2^H_7_]22S-HC constant. A plot of peak area ratio of [^2^H_7_]24R/S-HC: [^2^H_7_]22S-HC (averaged over three closely arranged spots from the same isocortical region on each slice) against areal density of sprayed-on [^2^H_7_]24R/S-HC, gave a straight line with R^2^ > 0.999 (SI Appendix, Figure S5A). When this experiment was repeated but by plotting the peak area ratio of [^2^H_7_]24R/S-HC: 24S-HC, where 24S-HC refers to the endogenous molecule, against areal density of sprayed-on [^2^H_7_]24R/S-HC, for closely arranged spots (n = 3) in the same region of interest, on six successive slices, the result was a straight line of R^2^> 0.994 (SI Appendix, Figure S5B & 5C). Assuming the concentration of endogenous 24S-HC does not vary across the six slices, or adjacent spots within the same region of interest, the results indicate that deuterated-surrogates can be used as a reliable internal standard for quantification of endogenous sterols.

To assess the reproducibility of the EADSA-µLESA-LC-MS technology, adjacent spots (n = 6) in the isocortical region of a single slice of mouse brain were analysed. The mean value of areal density of 24S-HC was determined to be 0.947 ± 0.078 ng/mm^2^ (mean ± SD) giving a % coefficient of variation (% CV) of 8%. Assuming that the endogenous concentration of 24S-HC did not vary across the six adjacent spots the reproducibility of the method is satisfactory. Reproducibility was further assessed by analysing nine regions of interest from four closely cut slices from a single brain. The % CV for areal density in each individual region varied from 12.4 % for the striatum where the areal density of 24S-HC was high (1.415 ± 0.176 ng/mm^2^, n = 4) to 46.3 % in the white matter of the cerebellum where the areal density of 24S-HC was low (0.191 ± 0.088 ng/mm^2^, n = 4, SI Appendix, Figure S6A). However, there is likely some contamination of white matter of the cerebellum by grey matter and vice versa, accounting for the high % CV in these regions. Other than for the cerebellum, the inter-slice reproducibility was deemed acceptable. Similar analysis for cholesterol with conventional LC-MS gave a value of 387.3 ± 14.6 ng/mm^2^ (% CV = 4%) for five adjacent spots in the isocortex of a single tissue slice.

### 2.2. Imaging Cholesterol Metabolism in Different Regions of Wild Type Mouse Brain

Although cholesterol has been imaged in adult rodent brain, its precursors and metabolites have not. To demonstrate the potential of our platform and gain insight into spatial cholesterol metabolism in brain, we quantitatively imaged the sterol distribution in nine regions of interest, i.e. isocortex, striatum, thalamus, hippocampus, midbrain, pons, medulla, cerebellum (grey matter) and cerebellum (white matter) using µLESA-LC-MS with conventional-flow and nano-LC-MS formats (Figure 4).

**Figure 4.**
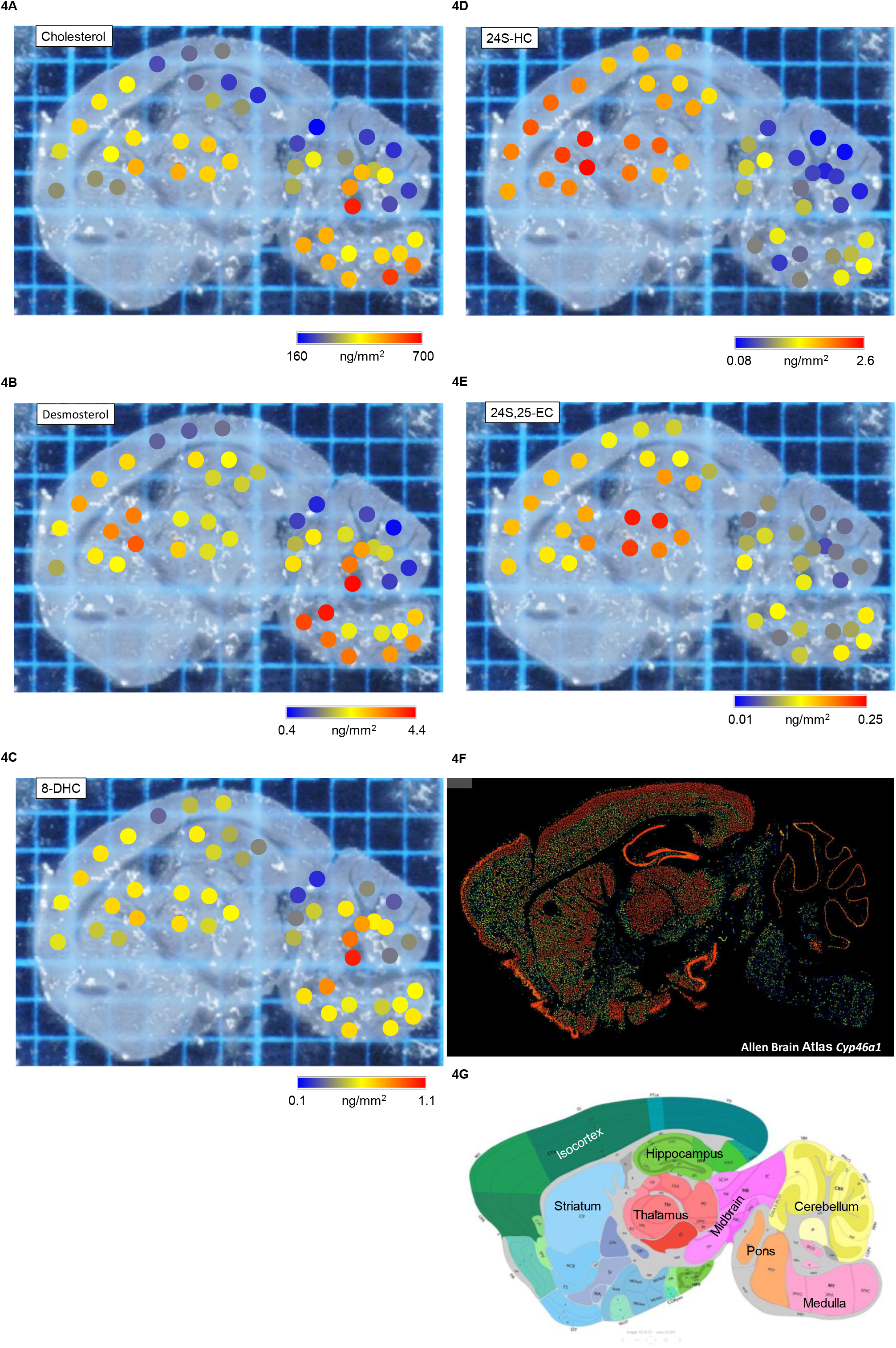
Sagittal section of a WT mouse brain analysed by µLESA-LC-MS following EADSA treatment. A photograph of a tissue slice showing spots analysed with areal density, determined by µLESA-LC-MS, of (A) cholesterol, (B) desmosterol, (C) 8-DHC, (D) 24S-HC and (E) 24S,25-EC. Areal density is colour coded on a blue to red scale as indicated. (F) Expression of *Cyp46a1* and (G) annotation of the different regions of mouse brain (57). Image credit for (F) and (G), Allen Institute. https://mouse.brain-map.org/experiment/show/532667 and http://mouse.brain-map.org/experiment/thumbnails/100042147?image_type=atlas.

Cholesterol is the most abundant sterol across the whole brain but there are concentration differences between regions (Figure 4A & 5A). Hierarchal cluster analysis separated the nine regions into three groups (Figure 5B), of which cholesterol is most abundant in pons (575.5 ± 122.0 ng/mm^2^, n = 3 animals) and white matter of the cerebellum (509.2 ± 83.0 ng/mm^2^) but least abundant in hippocampus (252.9 ± 27.3 ng/mm^2^), grey matter of cerebellum (263.3 ± 71.3 ng/mm^2^) and isocortex (305.1 ± 53.8 ng/mm^2^). Desmosterol, the immediate precursor of cholesterol in the Bloch pathway, was found to be about two orders of magnitude less abundant than cholesterol but its spatial distribution correlated significantly with cholesterol (SI Appendix, Figure S7A). The areal density of desmosterol ranged from 4.212 ± 1.546 ng/mm^2^ in pons to 1.025 ± 0.356 ng/mm^2^ in the grey matter of the cerebellum (Figure 4B & 5C). We also detected 8-DHC in our analysis (Figure 2B & 2D), which is an enzymatic isomeric product 7-DHC, the immediate precursor of cholesterol in the Kandutsch-Russell pathway (SI Appendix, Figure S1) (55). The areal density of 8-DHC was consistently lower than that of desmosterol in all brain regions measured, but its distribution pattern is quite similar to that of desmosterol and cholesterol (Figure 4 & 5). These data indicate that the distributions of cholesterol and its immediate precursors are correlated together in brain (SI Appendix, Figure S7A).

**Figure 5.**
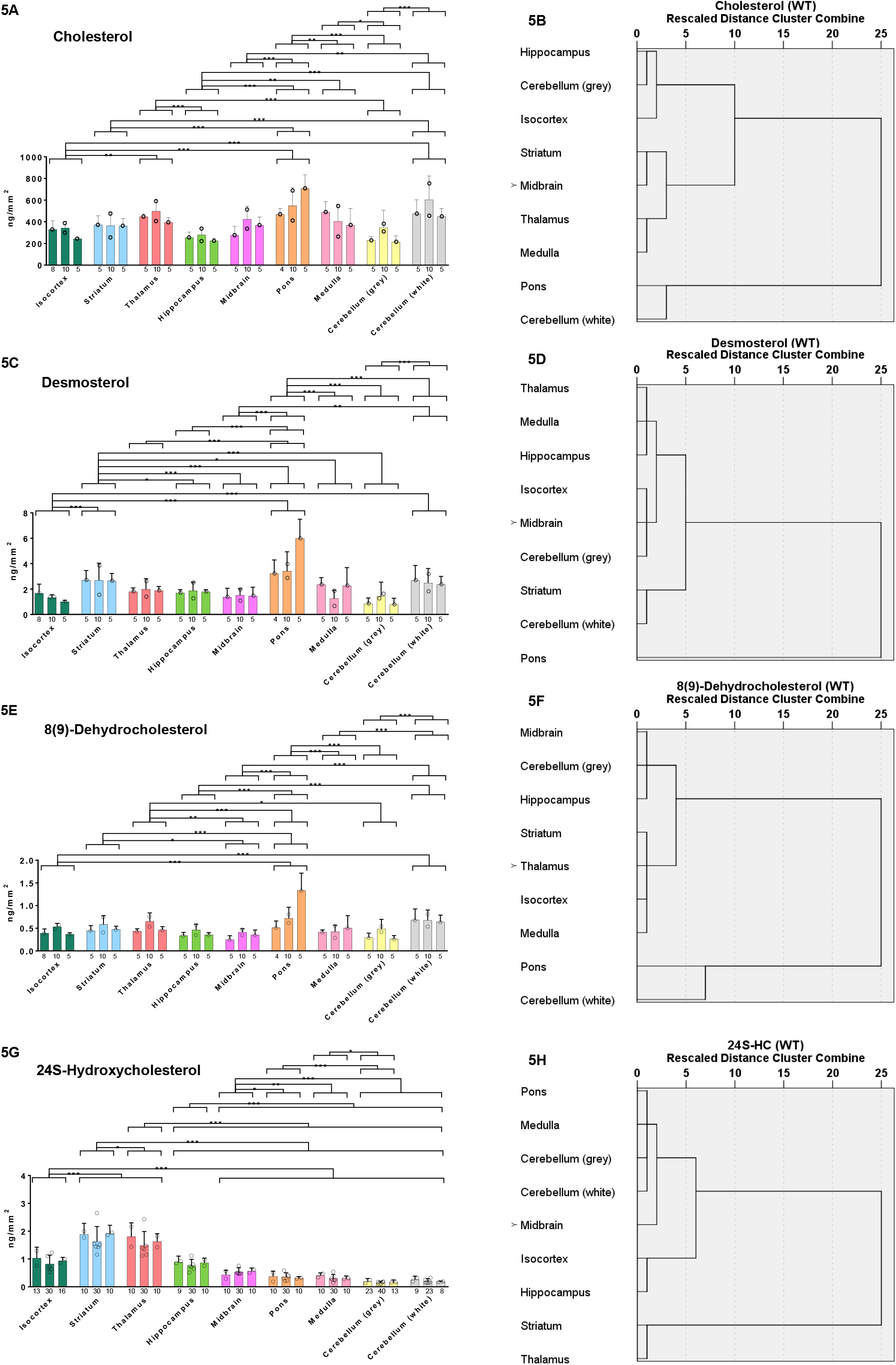
Sagittal section of WT mouse brain analysed by μLESA-LC-MS following EADSA treatment. Areal density (ng/mm^2^) of sterols in nine brain regions from three WT mice and dendrograms from hierarchal cluster analysis for the brain regions averaged over the biological replicates (see section 4.3). Areal density of (A) cholesterol, (C) desmosterol, (E) 8-DHC and (G) 24S-HC. Significance levels from ANOVA using Tukey’s multiple comparison test are indicated for those brain regions showing significant differences. *P*<0.05, *; *P*<0.01, **; *P*<0.001, ***. Values for individual mice are given by separate histogram bars. The number of dots within each bar indicates the number of brain slices analysed for each mouse. The dots within the bars correspond to region average for each brain slice. The height of each bar represents the mean of the region average of each brain slice. The number at the bottom of each bar indicates the total number of spots analysed per mouse. The error bars indicate the SD of all the spots per mouse. Dendrograms for (B) cholesterol, (D) desmosterol, (F) 8-DHC and (H) 24S-HC. A greater horizontal axis distance from zero indicates greater dissimilarity between different brain regions, whereas a lower distance from zero indicates greater similarity between different brain regions.

The major cholesterol metabolite in brain is 24S-HC, accounting for around 40% of cholesterol turn over in the brain. However, the distribution of 24S-HC is significantly different across different brain structures (Figure 4D & 5G), ranging in areal density from 0.161 ± 0.024 ng/mm^2^ in the grey matter of the cerebellum to 1.637 ± 0.159 ng/mm^2^ in the thalamus and 1.805 ± 0.158 ng/mm^2^ in the striatum. Remarkably, the distribution of 24S-HC across the different brain regions does not correlated significantly with levels of cholesterol or its precursors (SI Appendix, Figure S7A). Interestingly, while the level of 24S-HC varies by a factor of about ten across the different regions, the corresponding variations of cholesterol and desmosterol are only factors of about two and four, respectively. 24S-HC is formed from cholesterol by the enzyme CYP46A1 expressed in neurons localised to multiple sub-regions of brain (56). Interrogation of the Allen Mouse Brain Atlas reveals that *Cyp46a1* is highly expressed in striatum and thalamus but at lower levels in the cerebellum (Figure 4F) (57). Our results suggest that the pattern of 24S-HC in brain reflects local *Cyp46a1* expression.

Another oxysterol identified in brain is 24S,25-EC (Figure 2B & 2E) (15). 24S,25-EC is not a metabolite of cholesterol, but formed in parallel to cholesterol via a shunt pathway using the exact same enzymes, but with the exclusion of DHCR24 (SI Appendix, Figure S1) (14). In cellular systems, the level of 24S,25-EC reflects the activity of the cholesterol biosynthesis pathway. Recent studies also suggests that 24S,25-EC can be formed *in vitro* from desmosterol via a CYP46A1 catalysed reaction (16). We found 24S,25-EC to be present in all brain region but to be particular abundant in thalamus at an areal density of 0.158 ± 0.026 ng/mm^2^ (Figures 4E & 6A). Factor analysis reveals that 24S,25-EC correlates most closely with 24S-HC (SI Appendix, Figure S7A), although being about ten times less abundant. The distribution of 24S,25-EC in brain likely reflects the varying activities of *de novo* cholesterol biosynthesis and of CYP46A1 in the different brain regions.

While, cholesterol, desmosterol, 8-DHC, 24S-HC and 24S,25-EC can all be analysed by EADSA-µLESA-LC-MS at conventional flow rates, low abundance oxysterols require the more sensitive EADSA-µLESA-nano-LC-MS format. The nano-LC-MS format provides a lower limit of quantification (LOQ) of 0.01 ng/mm^2^ based on a CV of <25% at the LOQ and a limit of detection (LOD) of 0.001 ng/mm^2^ (SI Appendix, Figure S5).

Previous studies have suggested that 26-HC is imported into brain and converted in brain by a combination of enzymes, CYP27A1 and CYP7B1 to 3β,7α-diHCA and further by hydroxysteroid dehydrogenase (HSD) 3B7 to 7αH,3O-CA (SI Appendix, Figure S1) (21, 23, 24, 58). 3β,7α-diHCA is a ligand to the LXRs and is neuroprotective, while 7αH,3O-CA lacks these activities (21). Using EADSA-µLESA-nano-LC-MS 3β,7α-diHCA and 7αH,3O-CA are detected in brain. However, using the EADSA method 3β,7α-diHCA can be measured in combination with 7αH,3O-CA, and at low levels the combined areal densities of these two acids are measured more accurately than either of the individual acids alone. The combination of 3β,7α-diHCA plus 7αH,3O-CA was evident at areal densities ranging from close to the LOD i.e. 0.002 ± 0.002 ng/mm^2^ up to 0.018 ± 0.009 ng/mm^2^ across the different regions of interest in mouse brain (see SI Appendix, Table S1). The combined acids were found to be most abundant in the grey matter of the cerebellum, although not statistically different from the white matter of the cerebellum or isocortex (Figure 6G see also SI Appendix, Table S1).

**Figure 6.**
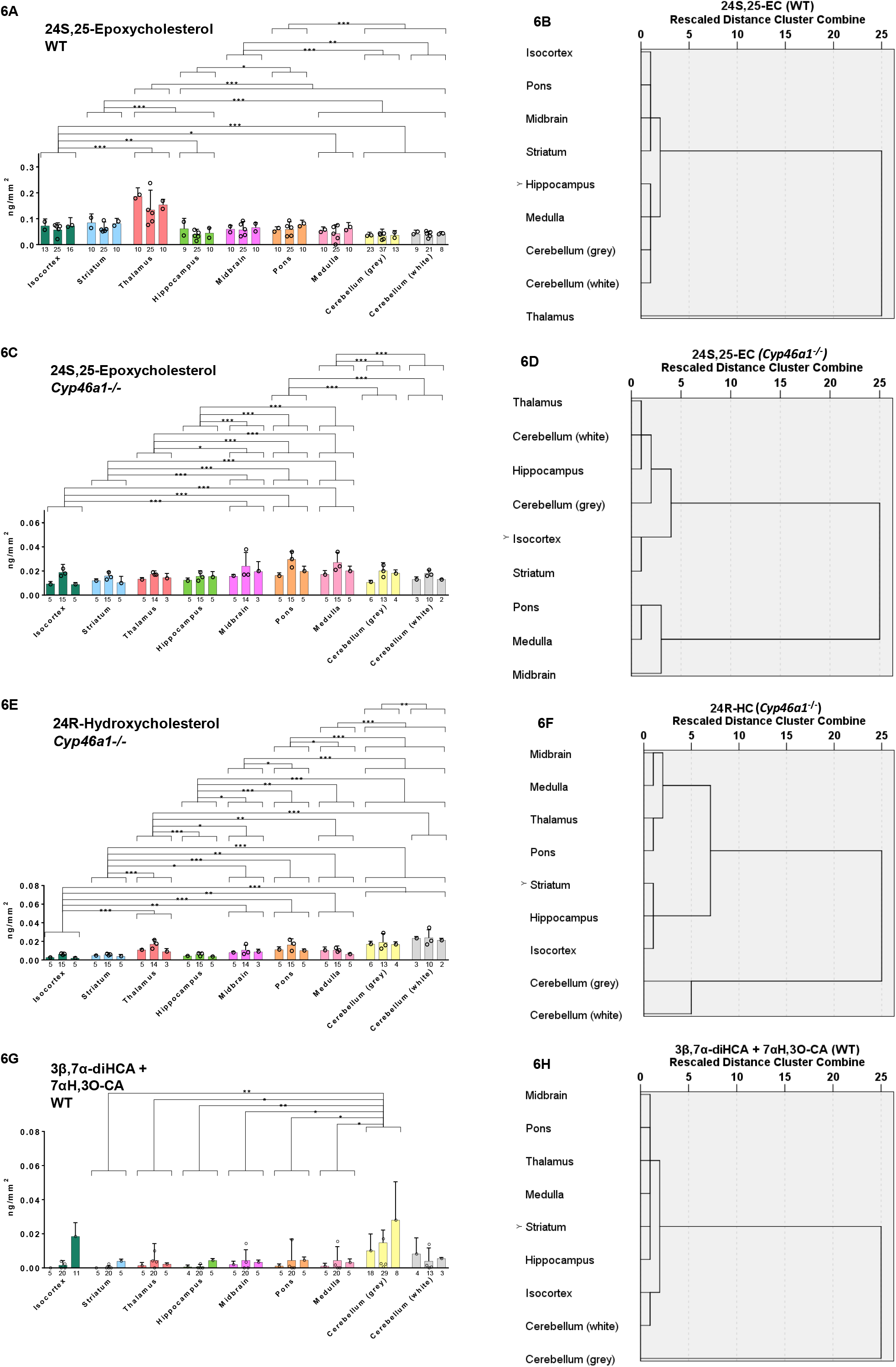
Sagittal section of WT and *Cyp46a1−/−* mouse brain analysed by μLESA-LC-MS following EADSA treatment. Areal density (ng/mm^2^) of sterols in nine brain regions from three mice of each genotype and dendrograms from hierarchal cluster analysis for the brain regions averaged over the biological replicates. Areal density of (A) 24S,25-EC, WT mouse; (C) 24S,25-EC, *Cyp46a1−/−* mouse; (E) 24R-HC, *Cyp46a1−/−* mouse; (G) combined 3β,7α-diHCA and 7αH,3O-CA, in WT mouse. Statistical analysis and labelling is as described in Figure 5. Dendrograms for (B) 24S,25-EC, WT mouse; (D) 24S,25-EC, *Cyp46a1−/−* mouse; (F) 24R-HC, *Cyp46a1−/−* mouse; (H) combined 3β,7α-diHCA and 7αH,3O-CA, in WT mouse.

7-OC can be formed from 7-DHC in a CYP7A1 catalysed reaction (59), or alternatively from cholesterol via free radical mediated reactions (60). As CYP7A1 is not expressed in brain, free radical formation or import across the BBB are the likely origins of this oxysterol in brain. 7-OC has previously been detected in brain using Girard derivatisation (37). We find it to be distributed ubiquitously across the brain at low levels (0.002 ± 0.002 – 0.004 ± 0.006 ng/mm^2^, see SI Appendix, Figure S8).

Taken together, using EADSA-µLESA-LC-MSI data we clearly demonstrate that different brain regions have distinct sterol patterns.

### 2.3. Imaging Biologically Active Oxysterols in Cyp46a1−/− Mouse Brain

Earlier studies have shown that although converting cholesterol to 24S-HC by CYP46A1 is responsible for the majority of cholesterol turnover in brain, the cholesterol content of whole brain does not differ significantly between wild type (WT) and *Cyp46a1−/−* animals (53, 54, 61). This was explained by a reduction of cholesterol synthesis compensating for reduced metabolism in the *Cyp46a1−/−* animals. As 24S,25-EC levels in brain may reflect the combined activities of *de novo* cholesterol synthesis and CYP46A1, we hypothesis that the levels of 24S,25-EC are reduced in *Cyp46a1−/−* mice. Indeed, we found that the level of 24S,25-EC decreased in all brain regions analysed. In particular, the areal densities of 24S,25-EC in the thalamus and striatum of *Cyp46a1−/−* animals were reduced by about 80 - 90% compared to WT brain (Figure 6A & 6C, see SI Appendix, Table S1), while there was only about a 50% reduction in medulla and cerebellum, demonstrating that a lack of CYP46A1 reduced 24S,25-EC in brain and that the thalamus and striatum are the most affected areas. Notably, the dendrograms from hierarchal cluster analysis are different in the two genotypes (Figure 6B & 6D).

In the WT mouse, the high areal density of 24S-HC (0.161 ± 0.024 - 1.805 ± 0.158 ng/mm^2^) compared to other oxysterols makes the identification of minor (≤ 0.01 ng/mm^2^) and closely eluting monohydroxycholesterols difficult. To overcome this problem, we have taken advantage of brain tissue from the *Cyp46a1−/−* mouse which is reported to be almost completely devoid to 24S-HC (53, 54, 61). Analysis of the same brain regions as for WT mouse confirmed an almost complete absence (≤0.001 ng/mm^2^) of 24S-HC in the *Cyp46a1−/−* animal (Figure 2C, see also SI Appendix, Table S1) but revealed minor amounts of 25-HC (0.005 ± 0.002 - 0.023 ± 0.034 ng/mm^2^) and 24R-HC (0.004 ± 0.002 – 0.023 ± 0.001 ng/mm^2^, SI Appendix, Figure S9). Although there was some variation in absolute concentrations in different mice, 24R-HC was at highest concentration in the cerebellum of each mouse (Figure 6E, see also SI Appendix, Table S1) and 25-HC was more ubiquitously dispersed (SI Appendix, Figure S8 and Table S1). Both of these oxysterols were detected at low levels in an earlier study of homogenised *Cyp46a1−/−* mouse brain, as was an unknown hydroxycholesterol eluting later than the peaks of 24R-HC (53). Here we again observe low levels of this oxysterol (0.007 ± 0.005 - 0.012 ± 0.006 ng/mm^2^) which we have previously assigned, based on retention time and MS^3^ spectra, but in the absence of an authentic standard, to be 12α-hydroxycholesterol (12α-HC, Figure S9A & S9E) (25).

Careful interrogation of the MS^3^ spectra recorded prior to the elution of 25-HC indicate the presence of not only trace quantities of 24S-HC (at about the limit of detection, SI Appendix, Figure S9A & S9C), but also of the elusive 20S-HC (SI Appendix, Figure S9B) (62). 20S-HC gives a unique MS^3^ spectrum where the fragment-ion *m/z* 327.2 is abundant (SI Appendix, Figure S9B & S9F). By generating a multiple reaction monitoring (MRM) chromatogram for the transition *m/z* 539.4→455.4→327.2 this oxysterol is highlighted (SI Appendix, Figure S9A). When the authentic standard of GP-derivatised 20S-HC was spotted on mouse brain tissue it eluted with an identical retention time to the GP-derivatised endogenous molecule and the MS^3^ spectra of the standard and endogenous molecule are almost identical. There are some small differences as a consequence of partial co-elution of isomeric oxysterols from tissue. However, an intense fragment-ion at *m/z* 327.2 is unique to 20S-HC. 20S-HC is a ligand of SMO and can activate the Hh signalling pathway and has previously been reported in only one prior publication, where it was found to be present in rat brain and human placenta (63). Here we define its presence in mouse brain. Our data demonstrate that 20S-HC is ubiquitously distributed in brain regions at an areal density of about 0.004 ± 0.003 to 0.007 ± 0.003 ng/mm^2^ in *Cyp46a1−/−* mice (SI Appendix, Figure S8 and Supplemental Table S1).

As discussed above, the combined areal densities of 3β,7α-diHCA and 7αH,3O-CA (SI Appendix, Figure S8) can be measured more accurately than either of the individual acids alone. In agreement with the data for WT mice, the combined acids were consistently most abundant in the grey matter of the cerebellum of the *Cyp46a1−/−* mice (0.013 ± 0.014 ng/mm^2^, SI Appendix, Figure S8), although there was some mouse to mouse variation in areal density.

Cholest-4-en-3-one has been found to be present in brain of Alzheimer’s disease sufferers and at lower levels in control brain samples (64). Here we detect cholest-4-en-3-one in all regions of mouse brain at areal densities in the region of 0.156 – 0.318 ng/mm^2^ in both WT and *Cyp46a1−/−* animals (SI Appendix, Figure S8).

### 2.4. Application of MALDI-MSI for Cholesterol Analysis

As an alternative to µLESA-LC-MS for the MSI of cholesterol we have explored the possibility of exploiting MALDI-MSI. Using on-tissue EADSA in a similar manner to that employed with µLESA, it was possible to image the [M]^+^ ion of GP-derivatised cholesterol (SI Appendix, Figure S10). There is good visual agreement between Figure S10 and Figure 5A.

## 3. Discussion

In the current work we have defined a methodology for the MSI of sterols including cholesterol, its immediate precursors and different oxysterols in mouse brain using EADSA and µLESA-LC-MS. This method compliments MALDI-MSI methods, also based on EADSA, being concurrently developed for cholesterol imaging in our laboratory. The µLESA-LC-MS method has a major advantage in that isomeric sterols and oxysterols can be separated by LC, even at the stereochemical level, something that has not been possible by any other separation devices linked to MS. µLESA-LC-MS technology can be operated at conventional LC flow-rates for rapid analysis or on a nano-LC-MS format to achieve high sensitivity for less abundant, but biologically important oxysterols. Note, that although an oxysterol may be minor in concentration, the metabolic pathway of which it is an intermediate may be significant. The major oxysterol in brain is 24S-HC (7, 65). Here we find 24S-HC to be most abundant in the striatum and thalamus and least abundant in the cerebellum. Unsurprisingly in the WT mouse, hierarchal cluster analysis for 24S-HC reveals that striatum and thalamus cluster together, as do grey and white matter of the cerebellum with pons and medulla (Figure 5H). 24S-HC is formed from cholesterol by the enzyme CYP46A1 expressed in neurons localised to multiple sub-regions of brain (56). *Cyp46a1*, is highly expressed in striatum and thalamus but at lower levels in the cerebellum (Figure 4F) (57), which reflects the abundance patterns of 24S-HC, and partially of 24S,25-EC, determined here by EADSA-µLESA-LC-MSI. The agreement between gene expression and enzyme product data evident here also indicates that 24S-HC does not rapidly move into other brain regions following its formation. In most studies, 24S-HC is analysed in whole brain homogenates being found at concentrations of 20 – 60 ng/mg (wet weight). Our values of 0.161 – 1.805 ng/mm (SI Appendix, Table S1) translate to 16 – 176 ng/mg assuming efficient extraction from the 10 µm thick tissue sections and minimal desiccation during tissue storage.

Cholesterol is the most abundant sterol in brain, being two to three orders of magnitude more abundant that 24S-HC. In the absence of derivatisation, or chelation with Ag^+^, cholesterol is barely detectable in most MSI experiments (29). However, following EADSA, cholesterol provides the most abundant signal in MSI analysis of brain tissue (Figure 2A). Cholesterol is found to be most abundant in the pons and white matter of the cerebellum (Figure 4A & 5A), unsurprisingly these two regions group together according to hierarchal cluster analysis (Figure 5B). Desmosterol, the immediate precursor of cholesterol in the Bloch pathway is most abundant in pons, as is 8-DHC, the isomer of 7-DHC the immediate precursor of cholesterol in the Kandutsch-Russell pathway (Figure 4B & C, Figure 5C & 5E). Factor analysis by sterol to sterol correlates cholesterol, desmosterol and 8-DHC most strongly (SI Appendix, Figure S7A). On the hand, 24S-HC correlates most strongly with 24S,25-EC. It is noteworthy that CYP46A1 generates 24S-HC from cholesterol and 24S,25-EC from desmosterol, providing one of the two pathways to biosynthesis of 24S,25-EC. Factor analysis clusters pons and white matter of the cerebellum together reflecting their high abundance of cholesterol and its precursors (SI Appendix, Figure S7B). Similarly, thalamus and striatum cluster together reflecting the abundance of 24S-HC and 24S,25-EC.

It is interesting to compare the distribution of sterols and the more abundant oxysterols with the expression of genes coding for their biosynthetic enzymes, as detailed in the Allen Mouse Brain Atlas (57). The Allen Mouse Brain Atlas (57) shows high expression in pons of 3-hydroxy-3-methylglutaryl-Coenzyme A synthase 1 (*Hmgcs1*), stearol-C-5 desaturase (*Sc5d*)*, Dhcr24* and dehydrocholesterol reductase 7 (*Dhcr7*), coding a key enzyme in the early part of the cholesterol biosynthesis pathway and three of the final enzymes in the pathway, respectively. There is also comparatively weak expression of *Cyp46a1*, coding cholesterol 24S-hydroxylase, in pons (57), which in combination can explain the elevated abundance of cholesterol in this region. According to the Allen Mouse Brain Atlas the expression of *Cyp27a1*, the gene coding the enzyme required to introduce a hydroxy and then a carboxylate group at C-26 of cholesterol, is of highest expression in the cerebellum (57), which is in agreement with data generated here, finding cholestenoic acids to be most abundant in cerebellum (Figure 6G).

24S-HC is by far the most abundant oxysterol in mouse brain (25, 53, 54). This makes analysis of closely eluting low abundance isomers challenging. However, deletion of *Cyp46a1* removes the biosynthesis of 24S-HC almost completely, and with implementation of nano-LC-MS, allows measurement of other isomers and oxysterols (53). In the current study we were able to image 24R-HC, the epimer of 24S-HC, which is found to be of most abundance in cerebellum (Figure 6E) and to detect other isomers including 12α-HC, 25-HC and the elusive 20S-HC (SI Appendix, Figure S8). 22R-HC was also found in some analysis but at the limit of detection. The enzymes required to biosynthesise 24R-HC and 20S-HC from cholesterol are unknown. The enhanced abundance of 24R-HC in cerebellum of the *Cyp46a1−/−* mouse does suggest CYP27A1 to be the relevant cholesterol hydroxylase, but 24R-HC is present in brain of the *Cyp27a1−/−* mouse arguing against this (25). CYP3A11 is a sterol 24R-hydroxylase and could perhaps convert cholesterol to 24R-HC in brain. *Cyp3a11* is however, predominantly expressed in cortex (57). CYP11A1 is the cholesterol 22R-hydroxylase required for neurosteroid biosynthesis and has been shown to be expressed in mouse brain (66). It first hydroxylates cholesterol at position C-22R, then at C-20, then cleaves the bond between these two carbons to generate pregnenolone. Besides being the enzyme responsible to generate 22R-HC from cholesterol, it may also be the catalyst for formation of 20S-HC from cholesterol, although evidence for this suggestion is lacking. There is only one previous report of the identification of 20S-HC in rodents or human (63). However, 20S-HC is of considerable biological interest being a ligand to the GPCR SMO, an integral receptor protein in the Hh signalling pathway, important for determining cell fate during development (67–69).

In terms of MSI, the spatial resolution and spot size available with the µLESA format (400 µm spot diameter), is admittedly inferior to that available with MALDI-MSI (<50 µm laser spot diameter). However, µLESA can be coupled with LC, offering isomer separation, essential for oxysterol analysis of tissue, where resolution of endogenous oxysterols from those generated *ex vivo* by oxidation in air is essential. MALDI can be linked with ion-mobility separation and MSI, but in experiments performed to-date we have been unable to separate different oxysterol isomers. A disadvantage with the µLESA-nano-LC-MS format adopted here is the length of chromatographic run time, this is however, reduced by increasing LC flow-rate and column diameter from a nano-scale to a more conventional scale (44, 46). However, increased flow-rate does come with the penalty of reduced sensitivity, due to the concentration dependency of ESI. This is not an issue for cholesterol, desmosterol, 8-DHC, 24S-HC or 24S,25-EC but is for less abundant oxysterols, where nano-LC-MS is more appropriate.

In summary, application of EADSA with µLESA-LC-MSI allows spatial location of oxysterols and cholestenoic acids in mouse brain. Through exploitation of the *Cyp46a1−/−* mouse, where the biosynthesis of 24S-HC is almost completely eliminated, other minor oxysterols are revealed. To compliment EADSA-µLESA-LC-MSI we are developing EADSA in combination with MALDI-MSI. We find this ideal for imaging cholesterol at high spatial resolution. EADSA-MSI, on a µLESA-LC-MSI or MALDI-MSI format, will be a powerful tool to study localised areas of healthy and diseased tissue.

## 4. Material and Methods

### 4.1. Experimental Design

The objective of the current study was to develop a MSI method suitable for the low-level quantitative determination of different sterols and oxysterols, including cholestenoic acids, in different regions of mouse brain.

#### 4.1.1. Chemicals and Reagents

HPLC grade methanol and water were from Fisher Scientific (Loughborough, UK). Glacial acetic and formic acids (FA) were purchased from VWR (Lutterworth, UK). [25,26,26,26,27,27,27-^2^H_7_]22S-Hydroxycholesterol ([^2^H_7_]22S-HC), [25,26,26,26,27,27,27-^2^H_7_]22R-hydroxycholesterol ([^2^H_7_]22R-HC), [25,26,26,26,27,27,27-^2^H_7_]24R/S-hydroxycholesterol ([^2^H_7_]24R/S-HC),[26,26,26,27,27,27-^2^H_6_]desmosterol and [25,26,26,26,27,27,27-^2^H_7_]cholesterol were from Avanti Polar Lipids (Alabaster, AL). [25,26,26,26,27,27,27-^2^H_7_]22S-Hydroxycholest-4-en-3-one ([^2^H_7_]22S-HCO) and [25,26,26,26,27,27,27-^2^H_7_]22R-hydroxycholest-4-en-3-one ([^2^H_7_]22R-HCO) were prepared from [^2^H_7_]22S-HC and [^2^H_7_]22R-HC, respectively, as described in Crick et al (46). Cholesterol oxidase from *Streptomyces sp.* and potassium dihydrogen phosphate were from Sigma-Aldrich, now Merck (Dorset, UK). [^2^H_0_]GP was from TCI Europe (Zwijndrecht, Belgium). [^2^H_5_]GP was synthesised as described by Crick et al (46, 70). Alpha-cyano-4-hydroxycinnamic acid (CHCA) was from Sigma-Aldrich (Dorset, UK).

#### 4.1.2. Tissue Sectioning

Mouse brain was from male 4-month-old *Cyp46a1−/−* and WT (*Cyp46a1+/+*, C57BL/6J;129S6/SvEv) littermates as detailed in (54). All animal experiments were approved by Case Western Reserve University’s Institutional Animal Care and Use Committee and conformed to recommendations made by the American Veterinary Association Panel on Euthanasia. Fresh frozen tissue, mounted in OCT (optimal cutting temperature compound), was cryo-sectioned using a Leica Cryostat CM1900 (Leica Microsystems, Milton Keynes, UK) at a chamber temperature of −16°C into 10 µm-thick sections which were thaw-mounted onto glass microscope slides and stored at −80°C until use. Luxol Fast Blue and Cresyl Violet staining was performed on tissue sections adjacent to sections analysed by MSI. For MALDI-MS, ITO (indium tin oxide) coated glass slides were from Diamond Coatings (Halesowen, UK).

#### 4.1.3. Deposition of Internal Standard and On-Tissue EADSA

Frozen tissue sections were dried in a vacuum desiccator for 15 min after which an internal standard mixture of [^2^H_7_]24R/S-HC (5 ng/µL), [^2^H_7_]22S-HC (1 ng/µL), [^2^H_7_]22S-HCO (1 ng/µL), in some experiments [^2^H_7_]22R-HCO (1 ng/µL), [^2^H_7_]cholesterol (20 ng/µL) and [^2^H_6_]desmosterol (5 ng/µL), in ethanol were sprayed from a SunCollect automated pneumatic sprayer (SunChrom, Friedrichsdorf, Germany supplied by KR Analytical Ltd, Cheshire, UK) at a flow rate of 20 µL/min at a linear velocity of 900 mm/min with 2 mm line distance and Z position of 30 mm in a series of 18 layers. The resulting density of the deuterated standard were, 1 ng/mm^2^ for [^2^H_7_]24R/S-HC, 0.2 ng/mm^2^ for each of [^2^H_7_]22S-HC, [^2^H_7_]22S-HCO and [^2^H_7_]22R-HCO, 4 ng/mm^2^ for [^2^H_7_]cholesterol and 1 ng/mm^2^ for [^2^H_6_]desmosterol (see SI Appendix). The sprayer was thoroughly flushed with about 2 mL of methanol after which cholesterol oxidase (0.264 units/mL in 50 mM KH_2_PO_4_, pH7) was sprayed for 18 layers. The first layer was applied at 10 µL/min, the second at 15 µL/min, then all the subsequent layers at 20 µL/min to give an enzyme density of 0.05 munits/mm^2^. Thereafter, the enzyme-coated slide was placed on a flat dry bed in a sealed pipette-tip box (11.9 cm x 8.2 cm x 8.5 cm) above 30 mL of warm water (37 °C), then incubated at 37°C for 1 hr. After which, the slide was removed and dried in a vacuum desiccator for 15 min. [^2^H_5_]GP (6.3 mg/mL bromide salt, in 70% methanol with 5% acetic acid) was sprayed on the dried slide with the same spray parameters as used for spraying cholesterol oxidase. The resulting GP density was 1.21 µg/mm^2^. The slide was then placed on a flat dry table in a covered glass chamber (12 cm x 12 cm x 7.2 cm) containing 30 mL of pre-warmed (37°C) 50% methanol, 5% acetic acid and incubated in a water bath at 37°C for 1 hr. The slide was removed and dried in a vacuum desiccator until LC-MS analysis. To analyse sterols containing a naturally occurring oxo group the cholesterol oxidase spray step was omitted and [^2^H_0_]GP (5 mg/mL chloride salt, in 70% methanol with 5% acetic acid) used as the derivatisation agent.

For MALDI-MSI experiments the above procedure was slightly modified as described in SI Appendix.

### 4.2. µLESA-LC-MSI

#### 4.2.1. Robotic µLESA

Derivatised oxysterols present in 10 µm thick slices of brain tissue were sampled using a TriVersa Nanomate (Advion, Ithaca, NY) with a modified capillary extraction probe (LESA^PLUS^). The Advion ChipsoftX software was used to capture an optical image of the prepared slide on a flatbed scanner prior to analysis. The same software was also used to define extraction points on the tissue prior to analysis. A 330 µm i.d. / 794 µm o.d. FEP sleeve was attached to a 200 µm i.d. / 360 µm o.d. fused silica capillary held in the mandrel of the Advion TriVersa Nanomate, this created a seal when in contact with tissue preventing the dispersion of extraction solvent (50% methanol) and limiting the sampling area to the internal diameter of the FEP sleeve (see SI Appendix, Figure S3A-B).

Three µL of extraction solvent was aspirated into the capillary from the solvent reservoir (R1 – R2 in Figure 3, SI Appendix, Figure S4), the mandrel moved robotically to an exact tissue location and 1 µL of solvent “dispensed” on-tissue and held for 30 s before being aspirated back into the capillary. The mandrel moved once again, and the extract was dispensed into a well of a 384-well plate. The process was repeated twice more with extract dispensed into the same well, before the combined extract was injected via the extraction probe into the sample loop of the LC system (see below). The entire procedure was then repeated on a new extraction point. Multiple points were analysed to build up an oxysterol image of brain tissue.

#### 4.2.2. LC-MS(MS^n^) at Conventional flow-rate

LC separations were performed on an Ultimate 3000 UHPLC system (Dionex, now Thermo Scientific, Hemel Hempstead, UK). The µLESA extract was delivered from the sample loop onto a trap column (Hypersil Gold C_18_ guard column, 3 µm particle size, 10 x 2.1 mm, Thermo Fisher Scientific) using a loading solvent of 20% methanol at a flow-rate of 50 µL/min. The eluent from the trap column containing unreacted GP reagent was delivered to waste (Figure 3). After 10 min the trap column was switched in-line with the analytical column (Hypersil Gold C_18_ column, 1.9 µm particle size, 50 x 2.1 mm, Thermo Fisher). At 11 min, 1 min after valve switching, a gradient was delivered by the binary micropump-2 at 200 µL/min. Mobile phase A consisted of 33.3% methanol, 16.7% acetonitrile, 0.1% formic acid and mobile phase B consisted of 63.3% methanol, 31.7% acetonitrile 0.1% formic acid. The proportion of B was initially 20 %, and was raised to 80% B over the next 7 min and maintained at 80% B for 4 min, before being reduced to 20% B in 0.1 min, just after the trap column switched off-line, and the analytical column was re-equilibrated for 3.9 min. After switching off line at 22 min, the trap column was independently washed with 10 µL of propan-2-ol injected from the Nanomate and delivered with mobile phase B at 200 µL/min for 6 min using micropump-1. At 26 min, the proportion of B washing the analytical column was raised from 20 % to 80% over 3 min and held at 80% for a further 5 min before dropping back down to 20% in 0.1 min and equilibrated for 4.9 min, ready for the next surface extraction, giving a total run time of 39 min. At 29 min the trap column was switched back in line with the analytical column and then out of line at 34 min to check for the presence of any sterols retained by the trap column. The gradients are illustrated in SI Appendix, Figure S3C-D.

For ESI-MS analysis the column eluent was directly infused into an Orbitrap Elite mass spectrometer (Thermo Scientific) using a H-ESI II probe. To avoid cross-contamination between samples a blank injection of 6 µL of 50% MeOH was performed with the same 39 min method between injections of tissue extracts.

For each injection five scan events were performed: one high resolution scan (120,000 full-width at half maximum height definition at *m/z* 400) in the Orbitrap analyser in parallel to four MS^3^ scan events in the linear ion-trap. Quantification was by isotope dilution using isotope-labelled standard spayed on-tissue. MS^3^ scans were for the transitions [M]^+^→[M-Py]^+^→, where Py is the pyridine group of the GP-derivative.

#### 4.2.3. Nano-LC-MS(MS^n^)

Nano-LC separations were performed on an Ultimate 3000 UHPLC system. The µLESA extract was delivered from the sample loop onto a trap column (HotSep Tracy C_8_, 5 µm 100 Å, 5 mm x 0.3 mm, Kinesis Ltd, Bedfordshire, UK) using a loading solvent of 50% methanol, 0.1% formic acid at a flow-rate of 5 µL/min. The eluent from the trap column containing unreacted GP reagent was delivered to waste (see SI Appendix, Figure S4). After 13 min the trap column was switched in-line with the analytical column (ACE C_18_, 3 µm 100 Å, 100 mm x 0.1 mm, Aberdeen, Scotland) and a gradient delivered by the binary nanopump at 0.6 µL/min. Mobile phase A consisted of 50% methanol, 0.1% formic acid and mobile phase B consisted of 95% methanol, 0.1% formic acid. At 13 min, the time of valve switching, the proportion of B was 40%, and was raised to 60% B over the next 5 min and maintained at 60% B for 5 min, then raised again to 95% B over 5 min and maintained at 95% B for a further 38 min before being reduced to 20% B in 1 min (run time 67 min) and the column re-equilibrated for 11 min. The trap-column was switched out of line at 67 min and re-equilibrated for 11 min with 50% methanol, 0.1% formic acid prior to the next injection (see SI Appendix, Figure S3E). The column eluent was directly infused into the Orbitrap Elite mass spectrometer (Thermo Scientific, Hemel Hempstead, UK) using the Nanomate chip-based nano-ESI (Advion, Ithaca, NY) set at between 1.7 kV and 2.0 kV chip voltage. The scan events were as described above. To avoid cross-contamination between samples a column wash with injection of 10 µL of 100% propan-2-ol and a fast gradient of 36 min was performed between injections of tissue extracts (see SI Appendix, Figure S3F). The propan-2-ol was delivered to the trap column which was washed for 5 min before being switched in-line with the analytical column, which during this period was washed independently by raising the proportion of B from 20% to 95%. With the two columns in-line the proportion of B was maintained at 95% B for 19 min after which it returned to 20% within 1 min. The trap column was then switched off-line and both columns equilibrated for a further 11 min before the next sample was injected. The gradients are illustrated in SI Appendix Figure S3E-F.

#### 4.2.4. Quantification

To achieve reliable quantitative measurements known amounts of isotope labelled internal standards [^2^H_7_]24R/S-HC, [^2^H_6_]desmosterol, [^2^H_7_]cholesterol and [^2^H_7_]22S-HCO were sprayed on-tissue prior to the EADSA process. This procedure corrects for variation in derivatisation efficiency, surface extraction, extraction area, injection volume, and MS response. Quantification was made from [M]^+^ ion signals in appropriate reconstructed ion chromatograms (RICs). Approximate quantification of 20S-HC was performed by utilising the MRM [M]^+^→[M-Py]^+^→327.2 transition and quantifying against the transition [M]+→[M-Py]+→353.3 for [^2^H_7_]24R/S-HC (see SI Appendix, S9F).

### 4.3. Statistics

For three WT mice, and separately for the three knockout mice, unreplicated two-way ANOVA was performed with sterol areal density as dependent variable and mouse and brain region as factors. The interaction between mouse and brain region was used as error variance. The residuals representing the interaction deviations were approximately normally distributed. Tukey’s multiple comparisons test was used to identify significant differences between brain regions. Hierarchical Cluster Analysis using between groups linkage was used to represent differences and similarities of average sterol areal densities between brain regions averaged across the three biological replicates for each type of mouse, and with the differences represented as squared Euclidean distances. The analyses were performed using IBM SPSS Statistics 22 (IBM Corp, Armonk, NY, USA) and GraphPad Prism 7.02 software (GraphPad Software Inc, CA, USA). *P*<0.05, *; *P*<0.01, **; *P*<0.001, ***. Whiskers on bar graphs represent one standard deviation (SD).

## Acknowledgements

This work was supported by the UK Biotechnology and Biological Sciences Research Council (BBSRC, grant numbers BB/I001735/1 and BB/N015932/1 to WJG, BB/L001942/1 to YW). RA holds a Sêr Cymru Fellowship supported by the Welsh Government and the European Regional Development Fund. Work in IP’s laboratory is supported by the National Institute of Health (grant number R01 GM062882). Members of the European Network for Oxysterol Research (ENOR, https://www.oxysterols.net/) are thanked for informative discussions. Dr Ruth Andrew is thanked for advice on performing on-tissue derivatisation and Drs Hanne Røberg-Larsen and Steven Wilson are thanked for information regarding nano-LC separations. Dr Mark Wyatt of the Swansea University Mass Spectrometry Facility is thanked for allowing access to MALDI-MS instrumentation.

## Author Contributions

WJG and YW designed research; EY performed research; WJG, YW, EY, DOS, OWH analysed data; RA, MB, NM, IP, JN, MRC, OWH contributed new reagents/analytical tools; WJG, YW and EY wrote the paper. All authors discussed and contributed to the manuscript. All authors approved the final manuscript.

