## Supplementary material for "Imaging Sterols and Oxysterols in Mouse Brain Reveals Distinct Spatial Cholesterol Metabolism": SI

**Supplementary Information**

### 1. Supplementary Results

#### *1.1. On-Tissue EADSA*

##### *1.1.1. Oxidation with Cholesterol Oxidase*

By analogy to the in-solution protocol (1, 2), the enzyme catalysed reaction was carried out with cholesterol oxidase sprayed onto tissue in $KH_2PO_4$ buffer followed by incubation in a humid chamber for 1 hr at 37 $^{o}$C. Profiling of the tissue by µLESA-nano-LC-MS showed no evidence of underivatized sterols, i.e. $[M+H]^+$, $[M+Na]^+$, $[M+K]^+$, $[M+H-(H_2O)_n]^+$, $[M+Na-(H_2O)_n]^+$ or $[M+K-(H_2O)_n]^+$, of endogenous cholesterol or 24S-HC, or of added deuterated-standards. In contrast the 3-oxo-4-ene products once derivatised with GP reagent gave intense $[M]^+$ ions.

##### *1.1.2. Derivatisation with GP Reagent*

It was necessary to incubate the GP-coated tissue in a humid atmosphere to achieve efficient derivatisation. Based on the in-solution protocol (1, 2), a derivatisation atmosphere above a methanol, water and acetic acid solvent was investigated. Based on µLESA-nano-LC-MS ion-current the solvent compositions of (i) 50% methanol, 5% acetic acid, and of (ii) 70% methanol, 5% acetic acid gave similarly efficient derivatisation, while (iii) 95% methanol, 5% acetic acid gave an inferior derivatisation atmosphere (Figure S2A). At (i) 50% methanol, 5% acetic acid and (ii) 70% methanol, 5% acetic acid the measured peak area ratios for endogenous 24S-HC to sprayed-on [$^2H_7$]22S-HC were indistinguishable for spots within the isocortex region (Figure S2B).

##### *1.1.3. Lateral Dispersion*

As both (i) 50% methanol, 5% acetic acid, and (ii) 70% methanol, 5% acetic acid gave similar derivatisation efficiency it was necessary to determine which was superior for minimising lateral dispersion. This was achieved by treating an entire brain slice (coronal section) by on-tissue EADSA, but only spraying [$^2H_7$]22S-HC on the left hemisphere. By measuring, using µLESA-nano-LC-MS, the [$^2H_7$]22S-HC/24S-HC peak area ratio at increasing distances from the centre line an indication of lateral dispersion could be made. Bearing in mind the µLESA extraction spot size is about 400 µm in diameter, the [$^2H_7$]22S-HC/24S-HC peak area ratio fell from 4.5 x $10^{-2}$ on the side sprayed with [$^2H_7$]22S-HC to 1.06 x $10^{-2}$ (23% “carry over”) for a spot centred at 500 µm from the central line on the non-sprayed side and to 0.1 x $10^{-2}$ (2% “carry over”) for a spot centred at 1 mm from the centre line when using a 50% methanol, 5% acetic acid solvent (Figure S2C). Note, the pons, medulla, and cerebellum, the regions which show the lowest 24S-HC levels, were avoided. The equivalent ratios at 70% methanol, 5% acetic acid were 4.8 x $10^{-2}$, 2 x $10^{-2}$ (500 µm, 43% "carry over") and 1.1 x $10^{-2}$ (1 mm, 23% "carry over"). In terms of lateral dispersion, the 50% methanol, 5% acetic acid solvent was superior and when considering that for a spot 500 µm from the centre line the exterior of the extraction circle was only 300 µm from this line, and 100 µm from an adjacent spot (on the centre line), a 23% "carry over" was deemed satisfactory.

##### *1.2. Oxysterols in Cyp46a1-/- Mouse Brain*

7α,(25R)26-Dihydroxycholesterol (7α,26-diHC, also called 7α,27-dihydroxycholesterol) and 7α,(25R)26-dihydroxycholest-4-en-3-one (7α,26-diHCO, also called 7α,27-dihydroxycholest-4-en-3-one) represent potential precursors of 7α-hydroxycholestenoic acids in brain (Figure S1). Although the current LC gradient was not optimised with these two oxysterols in mind, their presence in both WT and *Cyp46a1-/-* brain is defined by appropriate chromatographic peaks and by $MS^3$ spectra consistent with their structures (Figure S11A-B). However, the additional presence of 7α,24S-dihydroxy and 7α,25-dihydroxy isomers, and of the 7α,(25S)26-dihydroxy epimers, not separating from the 7α,(25R)26-dihydroxy isomer in the current system, cannot be ruled out. The levels of the dihydroxysterols are low in brain of both genotypes (0.001 ± 0.001 - 0.006 ± 0.011 ng/$mm^2$).

#### 2. Supplementary Methods

##### *2.1. Deposition of Internal Standards, Cholesterol Oxidase and GP-hydrazine for µLESA-LC-MSI*

###### *2.1.1. Internal Standards*

An internal standard mixture in ethanol of [$^2H_7$]24R/S-HC (5 ng/µL), [$^2H_7$]22S-HC (1 ng/µL), [$^2H_7$]22S-HCO (1 ng/µL), in some experiment [$^2H_7$]22R-HCO (1 ng/µL), and [$^2H_7$]cholesterol (20 ng/µL) and [$^2H_6$]desmosterol (5 ng/µL) was sprayed on-tissue using a SunCollect automated pneumatic sprayer at a flow rate of 0.02 mL/min, a spray speed of 900 mm/min, a line distance of 2 mm, for 18 layers.

Using eq. 1, the areal density on brain of [$^2H_7$]24R/S-HC was calculated to be 1 ng/$mm^2$, to be 0.2 ng/$mm^2$ for each of [$^2H_7$]22S-HC, [$^2H_7$]22S-HCO, and where applicable [$^2H_7$]22R-HCO, 4 ng/$mm^2$ for [$^2H_7$]cholesterol and 1 ng/$mm^2$ for [$^2H_6$]desmosterol.

Areal density (mg/$mm^2$) = [(no. layers) x (concentration, mg/mL) x flow rate (mL/min)] / [spray speed (mm/min) x line distance (mm)] (eq. 1)

###### *2.1.2. Cholesterol Oxidase*

The cholesterol oxidase activity was 0.264 U/mL in the sprayed solution, using eq.1 and similar spray parameters as for the internal standards (see section 4.1.3, main text), this translates to an areal density of 0.05 mU/$mm^2$.

###### *2.1.3. GP-hydrazine*

The concentration of the bromine salt of [$^2H_5$]GP sprayed on tissue was 6.3 mg/mL, using similar spray parameters as above (see section 4.1.3, main text) this translated to an areal density of 1.21 µg/$mm^2$ (5.1 nmol/$mm^2$). When the chloride salt of [$^2H_0$]GP was spayed on tissue the concentration was 5 mg/mL giving an areal density of 0.96 µg/$mm^2$ (5.1 nmol/$mm^2$).

##### *2.2. Deposition of Internal Standard, Cholesterol Oxidase and GP-hydrazine for MALDI-MSI*

For imaging cholesterol the concentration of [$^2H_7$]cholesterol spray solution was 50 ng/µL and sprayed as in section 2.1.1, resulting in a density of 10 ng/$mm^2$. The cholesterol oxidase was sprayed at the same concentration as in 2.1.2 (0.264 U/mL) but in 100 µM (instead of 50 mM) $KH_2PO_4$ at pH 7. After spraying cholesterol oxidase, the enzyme-coated slide was placed on a PTFE bed in a covered glass chamber (12 cm x 12 cm x 7.2 cm) containing 30 mL of warm water (37°C), then incubated at 37°C for 1 hr. Afterwards, the slide was removed and dried in a vacuum desiccator for 15 min. Subsequently [$^2H_0$]GP (5 mg/mL chloride salt, in 70% methanol with 5% acetic acid) was sprayed as in 2.1.3 above. The resulting GP density was 0.96 µg/mm$^2$. The slide was then placed on a PTFE bed in a covered glass chamber containing 10 mL of pre-warmed (37°C) 50% methanol, 5% acetic acid and incubated in a water bath at 37°C for 1 hr. The slide was dried in a vacuum desiccator, which was then placed in a cold room overnight. On the next day the desiccator was allowed to reach room temperature and then the slide removed for spraying with the CHCA MALDI matrix. CHCA was sprayed from a HTX TM-Sprayer (HTX Technologies, NC, USA) at 5 mg/mL in water:propan-2-ol:acetonitrile (3:4:3, v:v:v) at a flow rate of 80 µL/min and a linear velocity of 1200 mm/min, with 2 mm line distance and a criss-cross deposition method which alternates vertical and horizontal passes, for a total of 8, with an offset of 1 mm, resulting in a matrix density of 1.3 µg/mm$^2$. The sprayer nozzle was heated at 70 °C.

##### 3. MALDI-MSI

Following EADSA treatment, tissues slices were analysed on an UltrafleXtreme MALDI TOF/TOF MS (Bruker Daltonics, Bremen, Germany) equipped with a Smartbeam™ laser, emitting light at 355 nm, and operated in the reflectron mode and positive polarity. For MSI, each mass spectrum was automatically acquired using the auto-execute method in flexControl (Bruker Daltonics) software in the range of *m/z* 400 – 1000. Spatial resolution was set at 50 µm using flexImaging 4.1 software (Bruker Daltonics). Laser power was tuned to optimise signal to noise (S/N) ratio without distortions of the baseline at 80% of maximum with instrument settings of Global Offset at 5%, Attenuator Offset at 40%, Attenuator Range at 40% and with the main laser parameter set as "2_small" and operating at 2 kHz. These parameters result in a diameter of the laser spot of 50 µm, according to factory specifications and as verified by visual inspection with the instrument camera. Extraction voltages were as follows: IS1 20.00 – 19.92 kV, IS2 17.90 – 17.82 kV, Lens 8.50 – 8.53 kV, Rfl 21.10 – 20.99 kV, Rfl2 10.95 – 10.89 kV. Reflector gain was set at 3.0X. Pulsed Ion Extraction was timed at 160 ns. A gated ion suppression was applied up to 375 *m/z*. Each raster was sampled with 200 shots in 5 steps for a total of 1000 shots per raster. Total acquisition time was typically about 11.5 hr for a total of ~27000 positions and a file size of ~24000 MB. The MALDI instrument was calibrated using a mixture of phosphatidylcholine and lysophosphatidylcholine of known composition and having masses in the range of interest (Avanti Polar Lipids). After measurement, imaging spectra were re-calibrated using the batch process in flexAnalysis. Data were analysed and visualized using flexImaging 3.0 (Bruker Daltonics), and SCiLS Lab 2.0 (SCiLS, Bremen, Germany) without any processing step. Data were visualized using "window" (ISTD) normalization. Mass filters were chosen with a width of 0.5 Da.

##### Data Availability Statement

The data that support the findings are available from the corresponding author upon request. There are no restrictions on data availability.

##### Supplementary References to Allen Brain Atlas

http://mouse.brain-map.org/search/show?page_num=0&page_size=20&no_paging=false&exact_match=false&search_term=cyp46a1&search_type=gene

http://atlas.brain-map.org/atlas?atlas=2&plate=100883770#atlas=2&plate=100883770&resolution=13.96&x=7487.950032552084&y=3808.1727091471357&zoom=-3&structure=477

http://mouse.brain-map.org/search/show?page_num=0&page_size=20&no_paging=false&exact_match=false&search_term=cyp27a1&search_type=gene

http://mouse.brain-map.org/search/show?page_num=0&page_size=20&no_paging=false&exact_match=false&search_term=cyp3a11&search_type=gene

**Supplementary Figures**

Figure S1. Simplified view of the import, export and biosynthesis of oxysterols and sterols in brain. Enzymes are shown in blue. The Bloch pathway proceeds via desmosterol, the Kandustch-Russell pathway through 7-dehydrocholesterol and the shunt pathway in parallel to the Bloch pathway.

Figure S2. Incubation in a humid chamber above a solution of 50% methanol (MeOH), 5% acetic acid (HOAc) for 1 hr at 37°C provides an atmosphere for efficient GP-derivatisation with minimal lateral dispersion. (A) 50% methanol, 5% acetic acid and 70% methanol, 5% acetic acid are superior derivatisation solutions to 95% methanol, 5% acetic acid in terms of measured peak area (PA). (B) Quantification of 24S-HC in the isocortex region using either 50% methanol, 5% acetic acid or 70% methanol, 5% acetic acid gives indistinguishable values. The photographs on the right show the spots analysed in this study, coronal sections. (C) There is less lateral dispersion with 50% methanol, 5% acetic acid than with 70% methanol, 5% acetic acid. The peak area ratio [$^2H_7$]22S-HC : 24S-HC is shown on a heat-map scale. [$^2H_7$]22S-HC was sprayed only on the left-hemisphere. Arrows indicates a spot centred at 1 mm from the mid-line separating the two hemispheres. The mid-line is indicated by enhanced thickness of the vertical grid line.

Figure S3. Modification of the LESA$^{PLUS}$ configuration to provide a direct seal with the tissue surface preventing spreading of organic solvent. (A) FEP sleeve (orange colour) surrounding a fused silica capillary making a direct seal on a section of mouse brain tissue. (B) Microscope view of imprints made after LESA sampling of brain tissue. The inner circles define the extraction spot, the outer circles define the o.d. of the FEP sleeve. Gradient for the (C) trap and (D) analytical column at conventional flow-rates, see schematic in Figure 3, and (E) for the analytical column in the nano-LC configuration during the analysis run and (F) during the column wash phase.

Figure S4. Schematic representation of the of the µLESA system linked to nano-LC-MS. With connections between ports 1 and 6 and between 3 and 2, the 6-port injection valve is in the load position. Once loaded, connections are made between ports 4 and 3, and between 6 and 5. Sample is transported to the 10-port valve and with connections made between ports 8 and 9 and ports 2 and 3 (on the 10-port valve), sterols are trapped on the $C_8$ trap-column. Port 1 is then connected to 2, port 9 to 10 and sterols eluted from the trap column to the analytical nano-LC column.

Figure S5. Sprayed-on [$^2H_7$]24R/S-HC as an internal standard for oxysterol quantification. (A) Ten point plot of the peak area (PA) ratio of [$^2H_7$]24R/S-HC : [$^2H_7$]22S-HC against areal density of [$^2H_7$]24R/S-HC sprayed on-tissue, keeping the areal density of sprayed-on [$^2H_7$]22S-HC constant. (B) Ten point plot of the peak area ratio of [$^2H_7$]24R/S-HC : 24S-HC against areal density of [$^2H_7$]24R/S-HC sprayed on-tissue. In (A) and (B) data is for n = 3 spots in the isocortical region for each of the ten [$^2H_7$]24R/S-HC internal standard areal densities (1 slice per density) as illustrated in (C). In (A) and (B) error bars represent SD.

Figure S6. Sagittal section of WT mouse brain analysed by µLESA-nano-LC-MS following EADSA treatment. (A) Areal density of 24S-HC in four nearby sagittal slices cut from a single brain. (B) Areal density of 24S-HC in six nearby slices cut from three different brains. Slices were taken from the right hemisphere 1 – 2 mm away from the midsagittal line. The number of spots analysed for each region of each brain slice is indicated by numbers within the bars. % CV is shown above the bars for each region.

Figure S7. Factor analysis by sterol to sterol for WT mice. (A) Correlation matrix showing that 24S-HC correlates significantly with 24S,25-EC and that cholesterol, desmosterol and 8-DHC correlate significantly together. (B) Factor score plot with data for each mouse represented by different circles with different colours representing different regions of brain.

Figure S8. Areal density of different sterols, oxysterols and cholestenoic acids in nine different regions of mouse brain. Panels with a blue outline are from WT mice, panels with a red outline are from *Cyp46a1-/-* mice. The number of mice for each analyte is indicated by n. Error bars indicate SD.

Figure S9. µLESA-nano-LC-MS($MS^n$) analysis of the thamamus region of *Cyp46a1-/-* mouse brain following EADSA treatment. (A) Top panel, RIC for the $[M]^+$ ion of monohydroxycholesterols (539.4368 ± 10 ppm). 2nd panel, MRM chromatogram 539.4→455.4→327.2 characteristic of 20S-HC. 3rd panel, MRM chromatogram 539.4→455.4→353.3 characteristic of 24-HC. Bottom panel, MRM chromatogram 546.5→462.4→353.3 characteristic of $[^2H_7]$24-HC. (B) $MS^3$ ($[M]^+$→$[M-Py]^+$→) spectra of 20S-HC recorded at 27.42 and 28.20 min. (C) $MS^3$ ($[M]^+$→$[M-Py]^+$→) spectra of 24S-HC eluting at 27.85 min and of $[^2H_7]$24S-HC eluting at the same time. (D) $MS^3$ ($[M]^+$→$[M-Py]^+$→) spectra of 24R-HC eluting at 29.55 min and of $[^2H_7]$24R-HC eluting at 29.52 min. (E) $MS^3$ ($[M]^+$→$[M-Py]^+$→) spectra of 25-HC and 12α-HC recorded at 28.70 and 31.80 min. The identification of 12α-HC is based on retention time and $MS^3$ fragmentation pattern in the absence of an authentic standard. Authentic standards are available for the other oxysterols. (F) Fragment ions characteristic of 24-HC and 20S-HC.

Figure S10. MALDI-MSI following EADSA treatment of a sagittal section of WT brain. (A) Image of the $[M]^+$ ion of cholesterol. (B) Different regions of mouse brain (3). Image credit: Allen Institute. http://mouse.brain-map.org/experiment/thumbnails/100042147?image_type=atlas.

Figure S11. µLESA-nano-LC-MS($MS^n$) analysis of the cerebellum (grey matter) region of *Cyp46a1-/-* mouse brain following treatment with cholesterol oxidase. (A) Top panel, RIC for the $[M]^+$ ion of dihydroxysterols (555.4317 ± 10 ppm). Bottom panel, RIC for the $[M]^+$ ion of 3β,7α-diHCA plus 7αH,3O-CA (569.4110 ± 10 ppm). (B) $MS^3$ spectra recorded at 23.26 – 24.40 min and 24.40 – 24.97 min corresponding to 7α,25-diHC and 7α,26-diHC, respectively. (C) $MS^3$ spectra recorded at 24.32 and 25.59 min, and corresponding to 3β,7α-diHCA plus 7αH,3O-CA.

Table S1. Sterols in mouse brain.

Figure S1

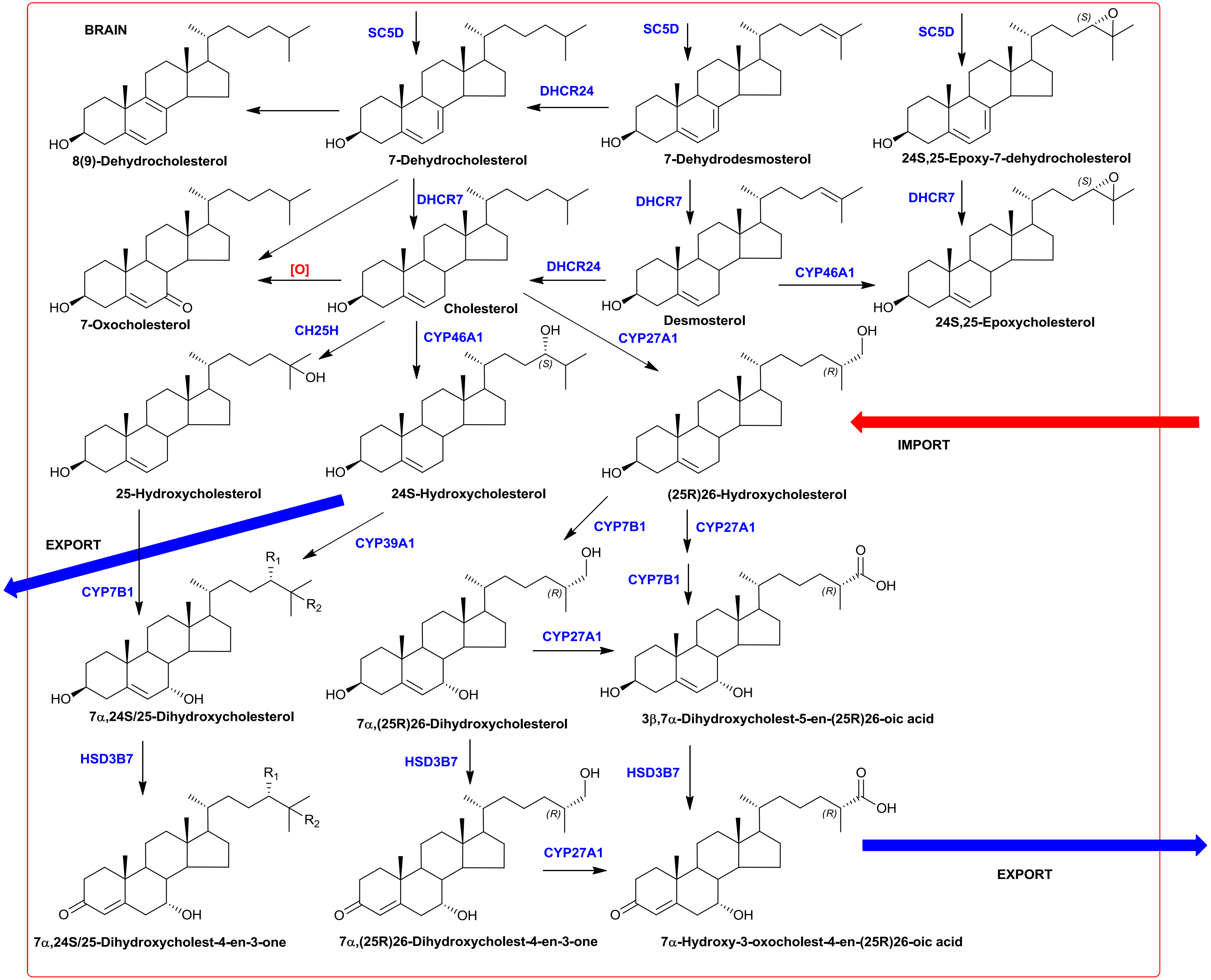

BRAIN
SC5D
DHCR24
DHCR7
CYP46A1
CYP27A1
CH25H
[O]
CYP7B1
CYP39A1
HSD3B7
IMPORT
EXPORT
8(9)-Dehydrocholesterol
7-Dehydrocholesterol
7-Dehydrodesmosterol
24S,25-Epoxy-7-dehydrocholesterol
7-Oxocholesterol
Cholesterol
Desmosterol
24S,25-Epoxycholesterol
25-Hydroxycholesterol
24S-Hydroxycholesterol
(25R)26-Hydroxycholesterol
7α,24S/25-Dihydroxycholesterol
7α,(25R)26-Dihydroxycholesterol
3β,7α-Dihydroxycholest-5-en-(25R)26-oic acid
7α,24S/25-Dihydroxycholest-4-en-3-one
7α,(25R)26-Dihydroxycholest-4-en-3-one
7α-Hydroxy-3-oxocholest-4-en-(25R)26-oic acid

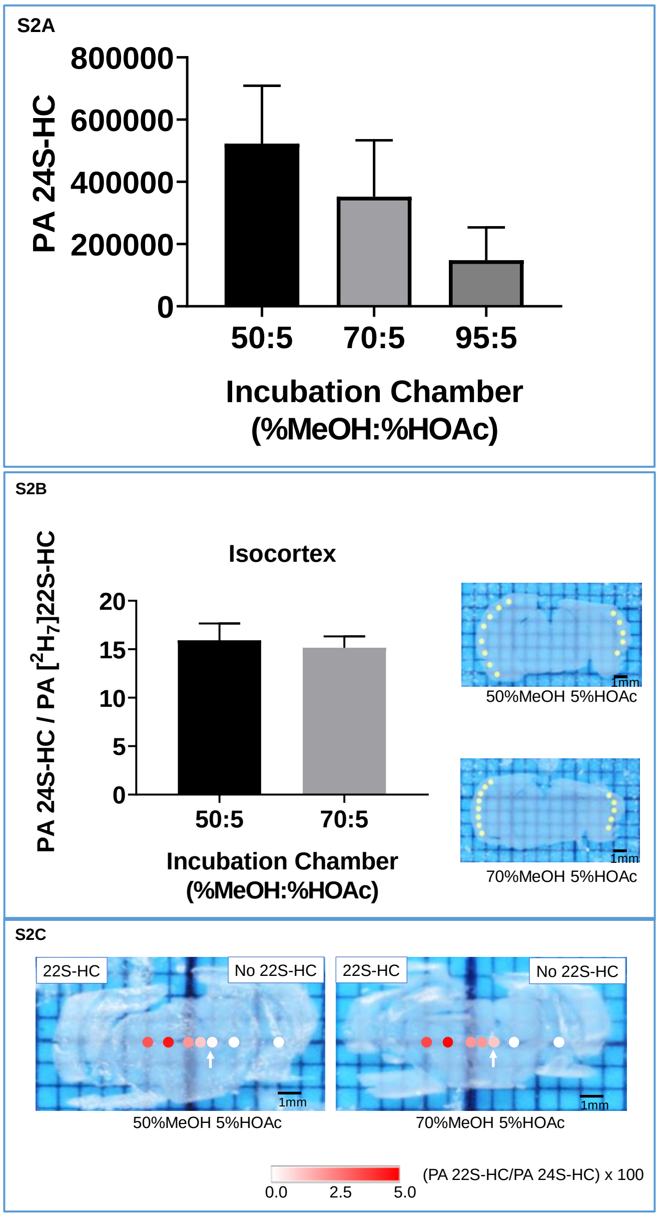
S2A
PA 24S-HC
800000
600000
400000
200000
0
50:5
70:5
95:5
Incubation Chamber
(%MeOH:%HOAc)
S2B
Isocortex
PA 24S-HC / PA [2H7]22S-HC
20
15
10
5
0
50:5
70:5
Incubation Chamber
(%MeOH:%HOAc)
1mm
50%MeOH 5%HOAc
1mm
70%MeOH 5%HOAc
S2C
22S-HC
No 22S-HC
1mm
50%MeOH 5%HOAc
22S-HC
No 22S-HC
1mm
70%MeOH 5%HOAc
0.0
2.5
5.0
(PA 22S-HC/PA 24S-HC) x 100

**S3A**

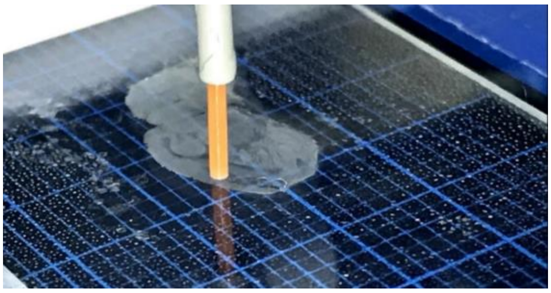

**S3B**

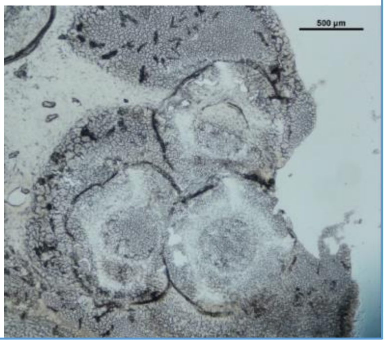


**S3C**

Trap Column Gradient

| min | %B |
|---|---|
| 0 | Loading solvent |
| 11 | 20 |
| 18 | 80 |
| 22 | 100 |
| 28 | 100 |
| 29 | 80 |
| 34 | Loading solvent |
| 39 | Loading solvent |

10 min. Trap column switched in-line with analytical column

22 min. Trap column switched off-line from analytical column

29 min.Trap column switched in-line with analytical column

34 min.Trap column switched off-line from analytical column

**S3D**

Analytical Column Gradient

| min | %B |
|---|---|
| 0 | 20 |
| 11 | 20 |
| 18 | 80 |
| 22 | 80 |
| 22.1 | 20 |
| 26 | 20 |
| 29 | 80 |
| 34 | 80 |
| 34.1 | 20 |
| 39 | 20 |

10 min. Trap column switched in-line with analytical column

22 min.Trap column switched off-line from analytical column

29 min. Trap column switched in-line with analytical column

34 min.Trap column switched off-line from analytical column

**S3E**

Analytical Column Gradient

| min | %B |
|---|---|
| 0 | 20 |
| 8 | 20 |
| 13 | 40 |
| 18 | 60 |
| 23 | 60 |
| 28 | 95 |
| 66 | 95 |
| 67 | 20 |
| 78 | 20 |

13 min
Trap column switched in-line with analytical column

67min
Trap column switched off-line from analytical column

**S3F**

Analytical Column Gradient

| min | %B |
|---|---|
| 0 | 20 |
| 5 | 95 |
| 24 | 95 |
| 25 | 20 |
| 36 | 20 |

5 min
Trap column switched in-line with analytical column

25 min
Trap column switched off-line from analytical column

S4

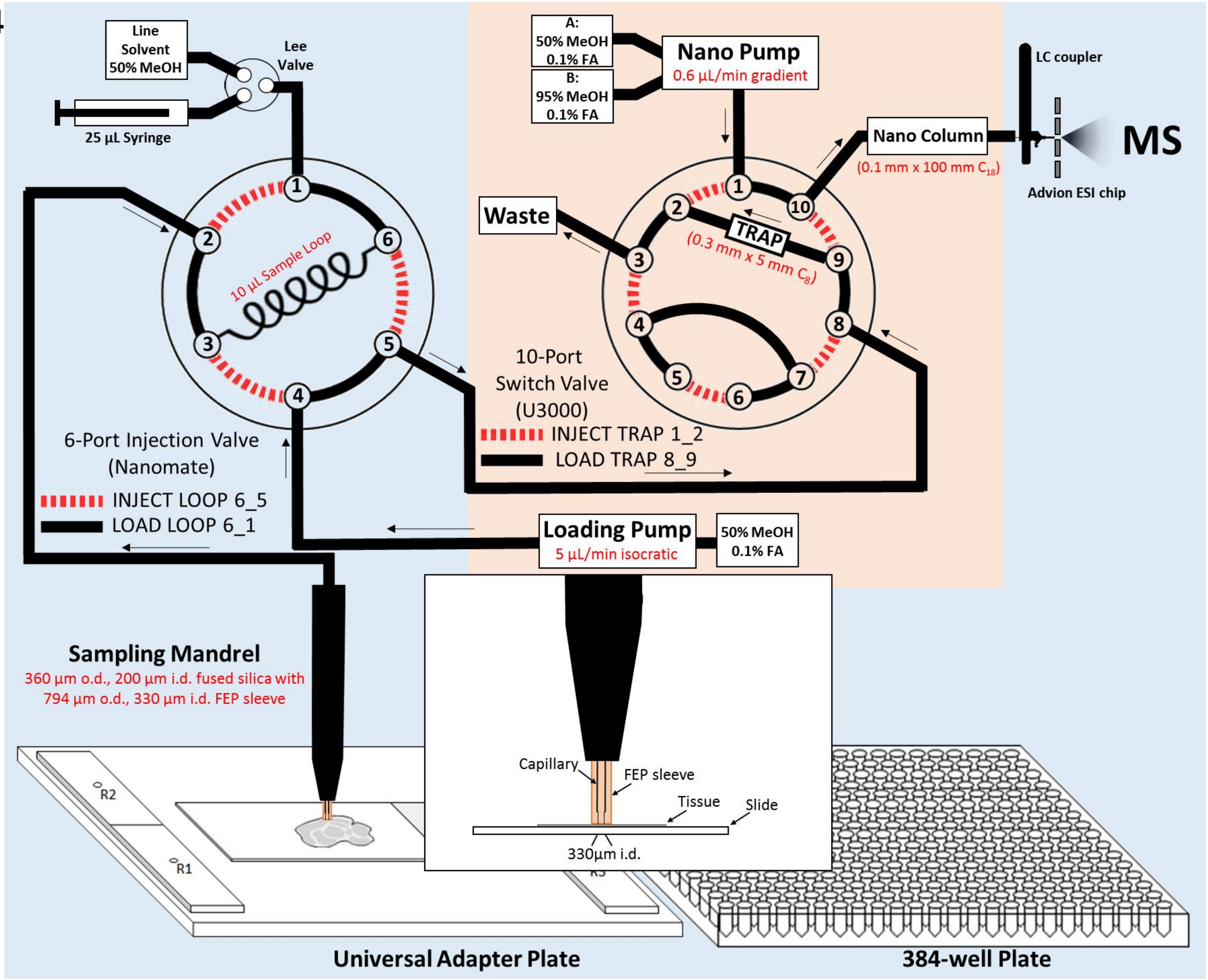
Line Solvent 50% MeOH
Lee Valve
25 µL Syringe
10 µL Sample Loop
6-Port Injection Valve (Nanomate)
INJECT LOOP 6_5
LOAD LOOP 6_1
A: 50% MeOH 0.1% FA
B: 95% MeOH 0.1% FA
Nano Pump
0.6 µL/min gradient
Waste
TRAP
(0.3 mm x 5 mm $C_8$)
10-Port Switch Valve (U3000)
INJECT TRAP 1_2
LOAD TRAP 8_9
Nano Column
(0.1 mm x 100 mm $C_{18}$)
LC coupler
MS
Advion ESI chip
Loading Pump
5 µL/min isocratic
50% MeOH 0.1% FA
Sampling Mandrel
360 µm o.d., 200 µm i.d. fused silica with 794 µm o.d., 330 µm i.d. FEP sleeve
Capillary
FEP sleeve
Tissue
Slide
330µm i.d.
R1
R2
Universal Adapter Plate
384-well Plate

S5A

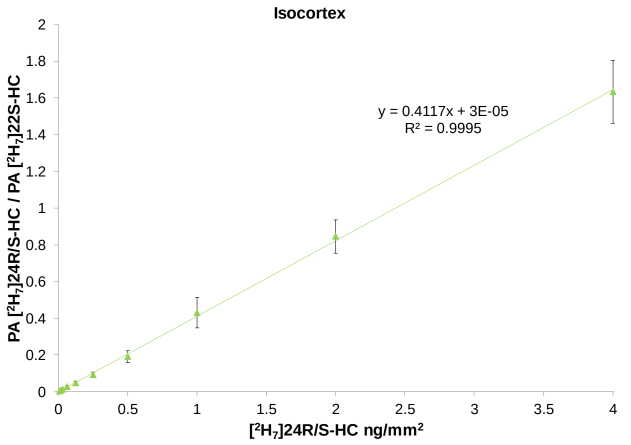
Isocortex
y = 0.4117x + 3E-05
R² = 0.9995
PA [2H7-]24R/S-HC / PA [2H7-]22S-HC
[2H7]24R/S-HC ng/mm2
0
0.2
0.4
0.6
0.8
1
1.2
1.4
1.6
1.8
2
0
0.5
1
1.5
2
2.5
3
3.5
4

S5B

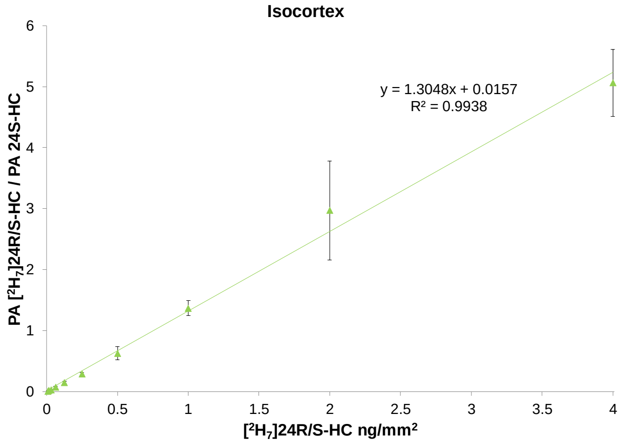
Isocortex
y = 1.3048x + 0.0157
R² = 0.9938
PA [2H7-]24R/S-HC / PA 24S-HC
[2H7]24R/S-HC ng/mm2
0
1
2
3
4
5
6
0
0.5
1
1.5
2
2.5
3
3.5
4

S5C

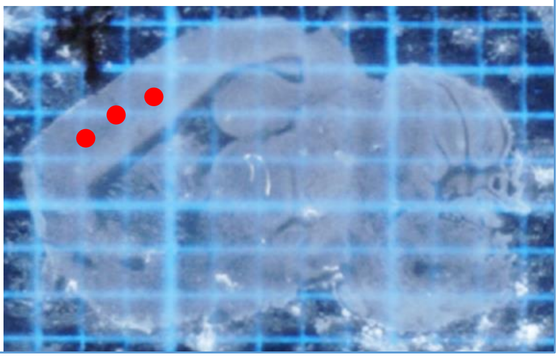

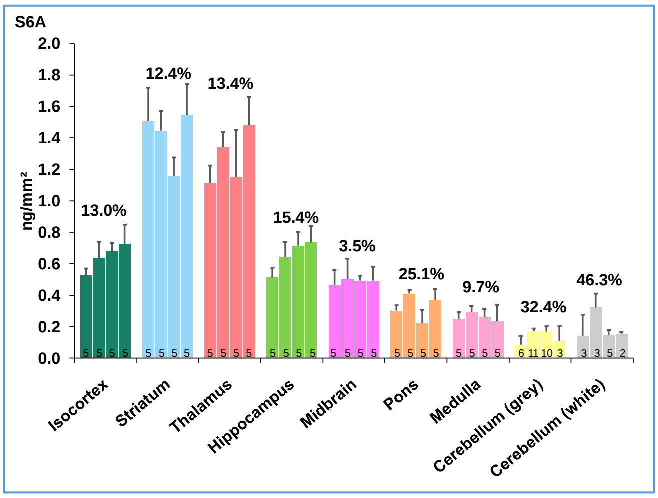
S6A
ng/mm²
2.0
1.8
1.6
1.4
1.2
1.0
0.8
0.6
0.4
0.2
0.0
13.0%
12.4%
13.4%
15.4%
3.5%
25.1%
9.7%
32.4%
46.3%
Isocortex
Striatum
Thalamus
Hippocampus
Midbrain
Pons
Medulla
Cerebellum (grey)
Cerebellum (white)

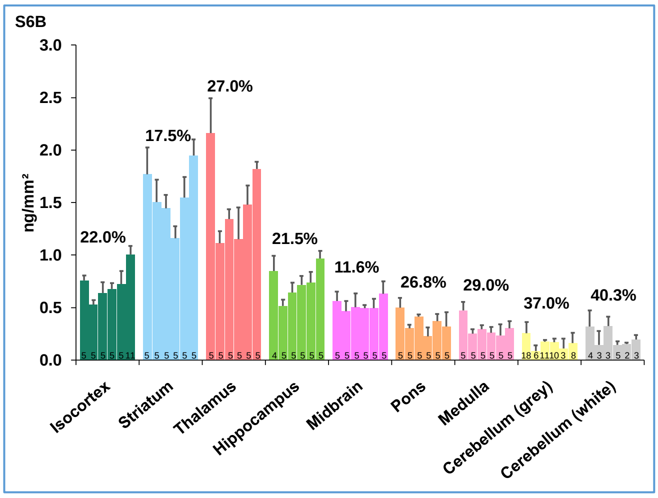
S6B
ng/mm²
3.0
2.5
2.0
1.5
1.0
0.5
0.0
22.0%
17.5%
27.0%
21.5%
11.6%
26.8%
29.0%
37.0%
40.3%
Isocortex
Striatum
Thalamus
Hippocampus
Midbrain
Pons
Medulla
Cerebellum (grey)
Cerebellum (white)

**S7A**

**Correlation Matrix**

| | | 24S-HC | 24S,25-EC | Cholesterol | Desmosterol | 8-DHC |
|---|---|---|---|---|---|---|
| Correlation | 24S-HC | 1.000 | .728 | -.118 | .004 | -.114 |
| | 24S,25-EC | .728 | 1.000 | .197 | .082 | .068 |
| | Cholesterol | -.118 | .197 | 1.000 | .776 | .828 |
| | Desmosterol | .004 | .082 | .776 | 1.000 | .884 |
| | 8-DHC | -.114 | .068 | .828 | .884 | 1.000 |
| Sig. (1-tailed) | 24S-HC | | .000 | .278 | .493 | .285 |
| | 24S,25-EC | .000 | | .162 | .342 | .368 |
| | Cholesterol | .278 | .162 | | .000 | .000 |
| | Desmosterol | .493 | .342 | .000 | | .000 |
| | 8-DHC | .285 | .368 | .000 | .000 | |

**S7B**

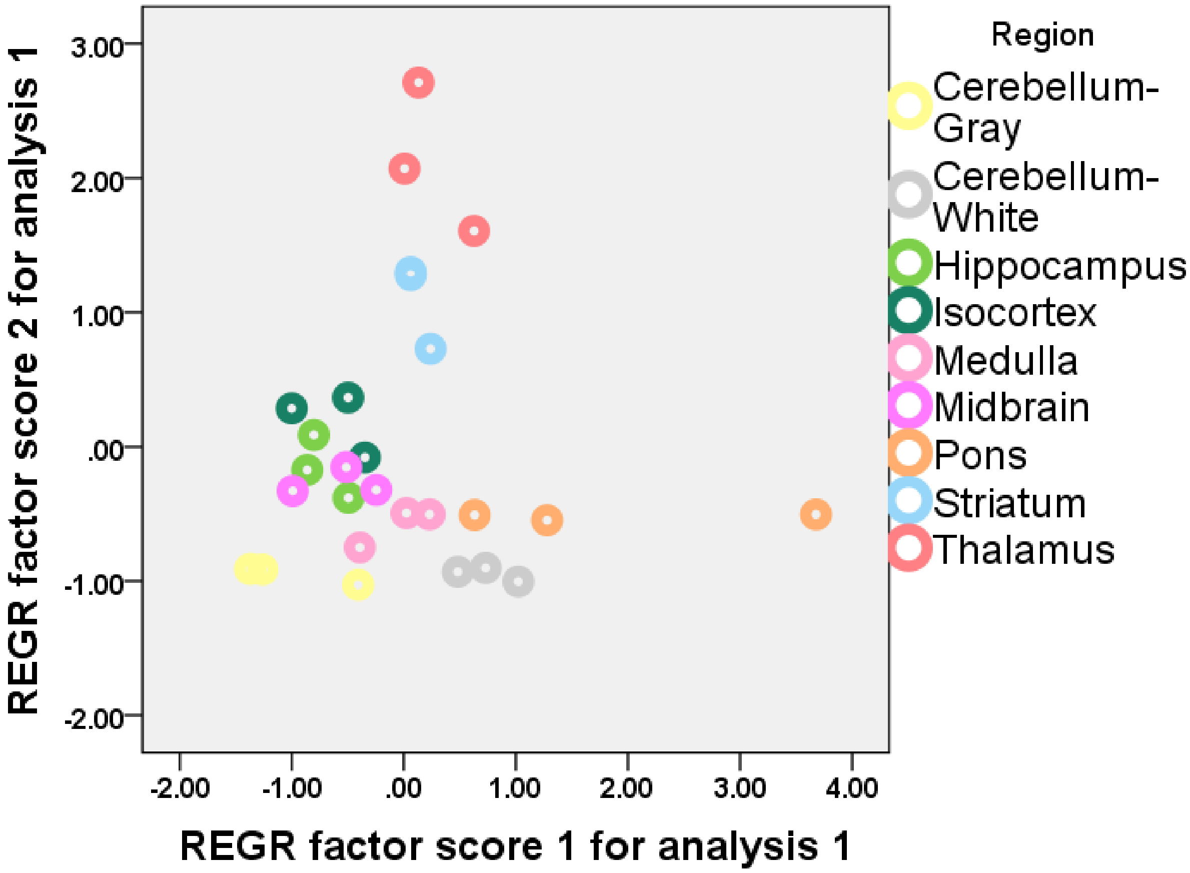

**S8**

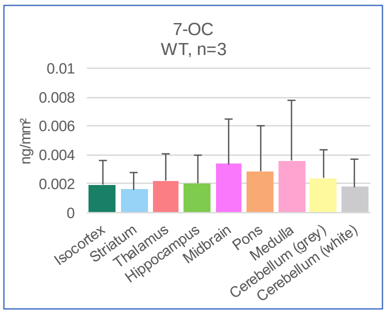


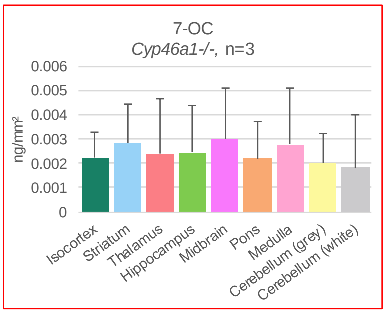


WT
Not analysed

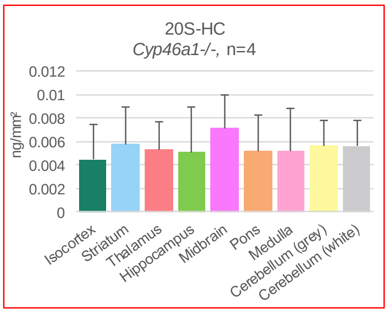


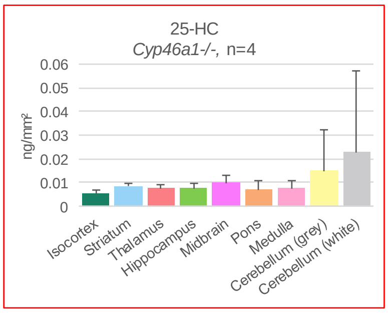


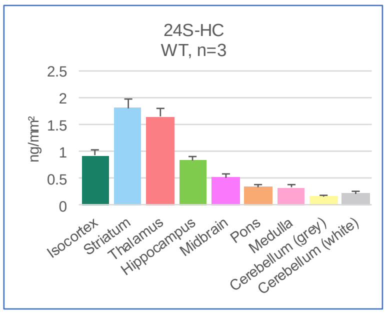


*Cyp46a1-/-*
Detected, not quantified

WT
Detected, not quantified

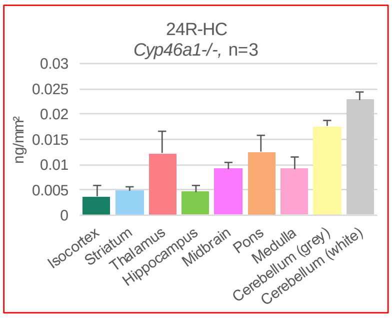


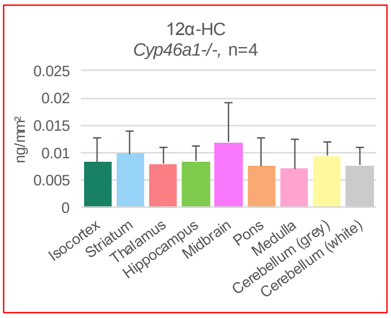


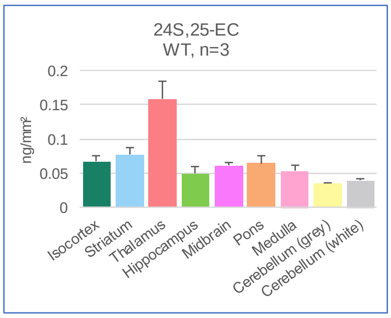


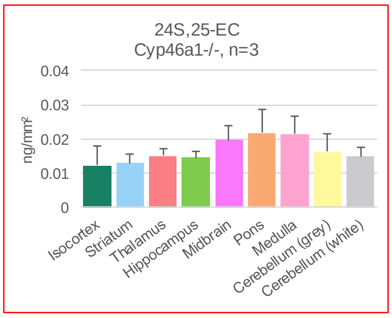


WT
Not analysed

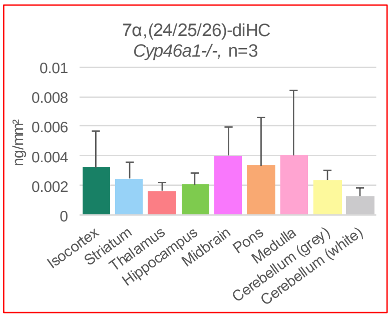

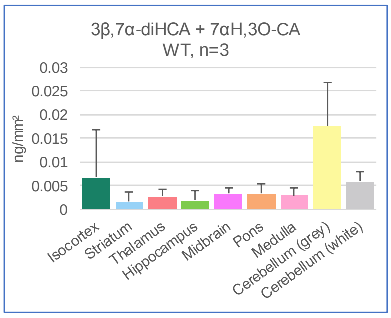


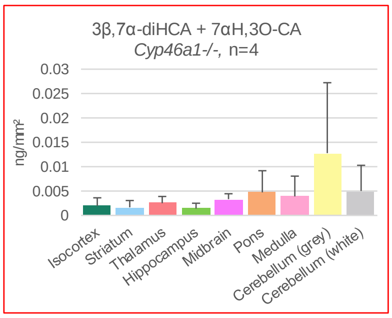


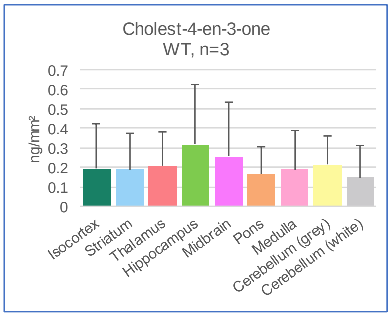


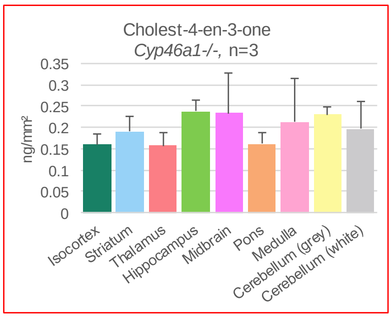


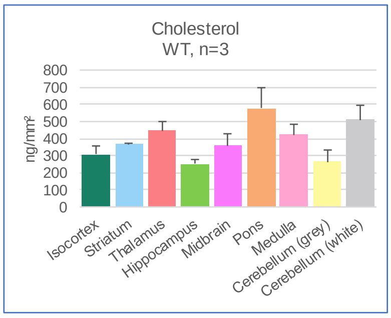


*Cyp46a1-/-*
Not analysed

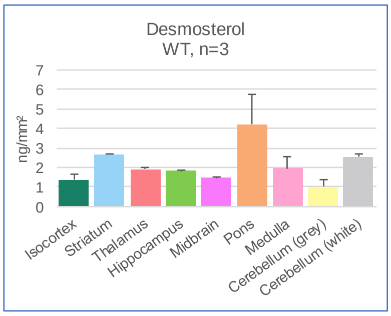


*Cyp46a1-/-*
Not analysed

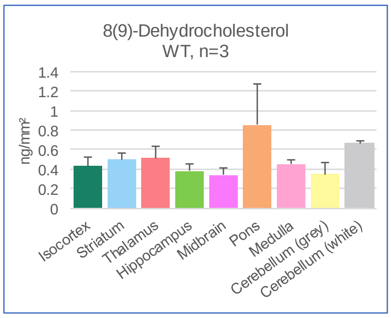


*Cyp46a1-/-*
Not analysed

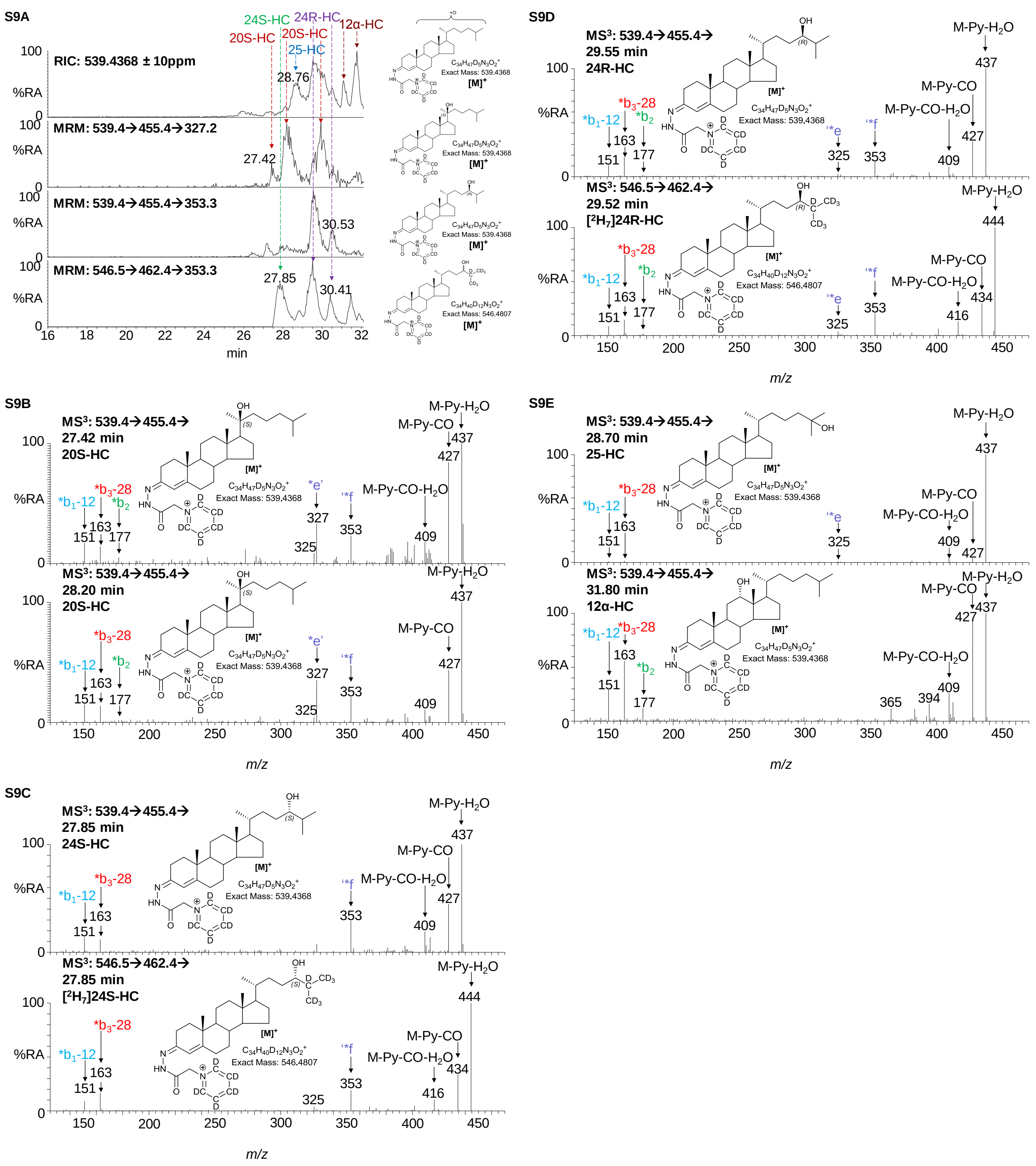
S9A
RIC: 539.4368 ± 10ppm
MRM: 539.4→455.4→327.2
MRM: 539.4→455.4→353.3
MRM: 546.5→462.4→353.3
24S-HC
24R-HC
20S-HC
25-HC
12α-HC
28.76
27.42
30.53
27.85
30.41
%RA
min
$C_{34}H_{47}D_5N_3O_2^+$
Exact Mass: 539.4368
$C_{34}H_{40}D_{12}N_3O_2^+$
Exact Mass: 546.4807
[M]+
S9B
MS3: 539.4→455.4→
27.42 min
20S-HC
28.20 min
S9C
27.85 min
24S-HC
MS3: 546.5→462.4→
[2H7]24S-HC
S9D
29.55 min
24R-HC
29.52 min
[2H7]24R-HC
S9E
28.70 min
25-HC
31.80 min
12α-HC
M-Py-H2O
M-Py-CO
M-Py-CO-H2O
*b1-12
*b3-28
*b2
*e'
*f
*e
m/z

**S10**

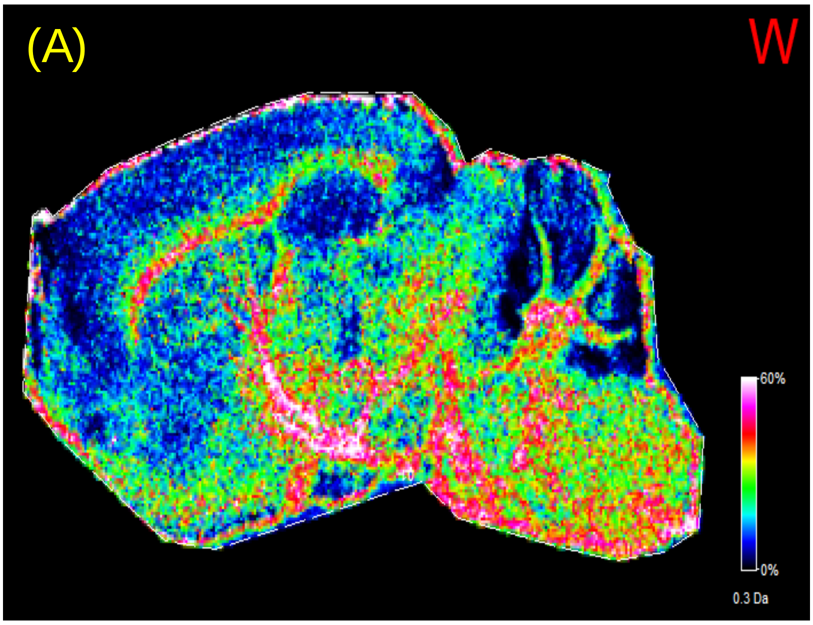
(A)
W
60%
0%
0.3 Da

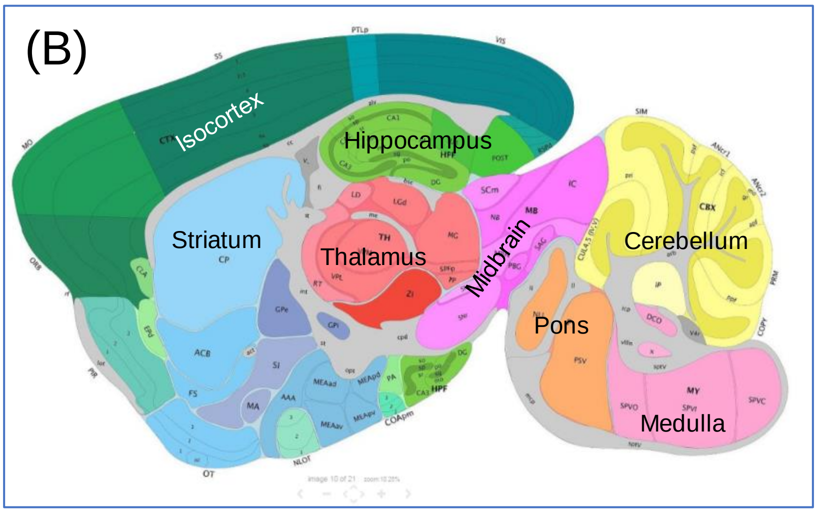
(B)
Isocortex
Hippocampus
Striatum
Thalamus
Midbrain
Cerebellum
Pons
Medulla

S11A

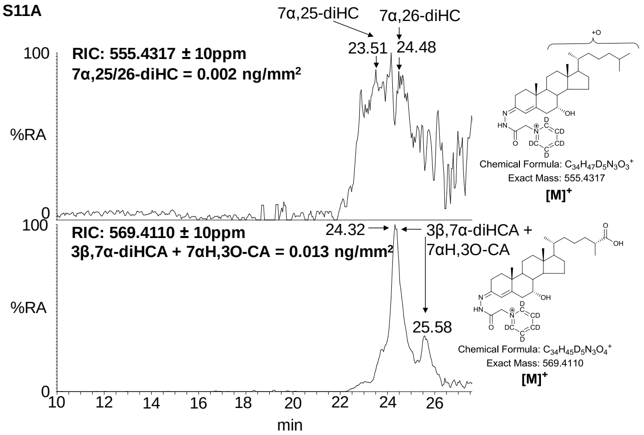


S11B

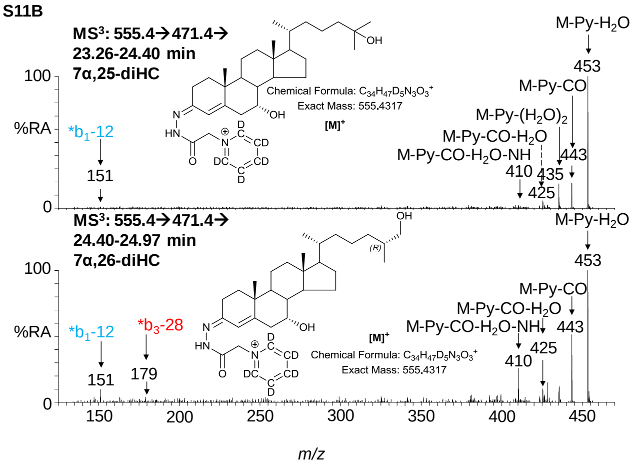


S11C

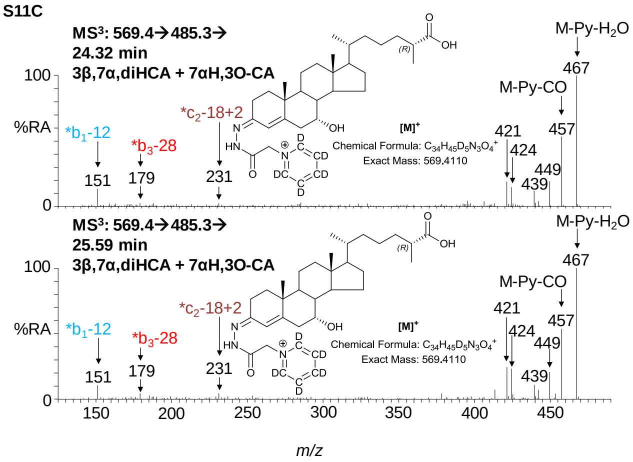
