## Supplementary material for "Imaging Sterols and Oxysterols in Mouse Brain Reveals Distinct Spatial Cholesterol Metabolism": SI Table

**Table S1. Sterols in mouse brain**

| LipidMaps ID | Systematic (common name) | Systematic abbreviation | Common abbreviation | Mouse type | | Sterols in mouse brain (ng/mm²) Isocortex | Striatum | Thalamus | Hippocampus | Midbrain | Pons | Medulla | Cerebellum (grey) | Cerebellum (white) | Note |
|---|---|---|---|---|---|---|---|---|---|---|---|---|---|---|---|
| LMST01010012 | 3β-Hydroxycholest-5-en-24S,25-epoxide (24S,25-Epoxycholesterol) | $C^{5}$-3β-ol-24S,25-epoxide | 24S,25-EC | WT (n=3) | Mean | 0.067 | 0.076 | 0.158 | 0.049 | 0.061 | 0.064 | 0.053 | 0.035 | 0.039 | 1 |
| | | | | | SD | 0.009 | 0.011 | 0.026 | 0.011 | 0.004 | 0.011 | 0.010 | 0.001 | 0.002 | |
| | | | | $Cyp46a1^{-/-}$ (n=3) | Mean | 0.012 | 0.013 | 0.015 | 0.014 | 0.020 | 0.022 | 0.021 | 0.016 | 0.015 | |
| | | | | | SD | 0.006 | 0.003 | 0.002 | 0.002 | 0.004 | 0.007 | 0.005 | 0.005 | 0.003 | |
| No ID | Cholest-5-ene-3β,12α-diol (12α-Hydroxycholesterol) | $C^{5}$-3β,12α-diol | 12α-HC | WT (n=0) | Mean | NA | NA | NA | NA | NA | NA | NA | NA | NA | 2 |
| | | | | | SD | | | | | | | | | | |
| | | | | $Cyp46a1^{-/-}$ (n=4) | Mean | 0.008 | 0.010 | 0.008 | 0.008 | 0.012 | 0.008 | 0.007 | 0.009 | 0.008 | 3 |
| | | | | | SD | 0.004 | 0.003 | 0.002 | 0.003 | 0.006 | 0.004 | 0.005 | 0.002 | 0.003 | |
| LMST01010134 | Cholest-5-ene-3β,20S-diol (20S-Hydroxycholesterol) | $C^{5}$-3β,20S-diol | 20S-HC | WT (n=0) | Mean | NA | NA | NA | NA | NA | NA | NA | NA | NA | 2 |
| | | | | | SD | | | | | | | | | | |
| | | | | $Cyp46a1^{-/-}$ (n=4) | Mean | 0.004 | 0.006 | 0.005 | 0.005 | 0.007 | 0.005 | 0.005 | 0.006 | 0.006 | |
| | | | | | SD | 0.003 | 0.003 | 0.002 | 0.004 | 0.003 | 0.003 | 0.004 | 0.002 | 0.002 | 4 |
| LMST01010086 | Cholest-5-ene-3β,22R-diol (22R-Hydroxycholesterol) | $C^{5}$-3β,22R-diol | 22R-HC | WT (n=0) | Mean | NA | NA | NA | NA | NA | NA | NA | NA | NA | 2 |
| | | | | | SD | | | | | | | | | | |
| | | | | $Cyp46a1^{-/-}$ (n=3) | Mean | LLD | LLD | LLD | LLD | LLD | LLD | LLD | LLD | LLD | 5 |
| | | | | | SD | 0.000 | 0.000 | 0.000 | 0.000 | 0.000 | 0.000 | 0.000 | 0.000 | 0.000 | |
| LMST01010019 | Cholest-5-ene-3β,24S-diol (24S-Hydroxycholesterol) | $C^{5}$-3β,24S-diol | 24S-HC | WT (n=3) | Mean | 0.926 | 1.805 | 1.637 | 0.834 | 0.507 | 0.342 | 0.329 | 0.161 | 0.211 | |
| | | | | | SD | 0.105 | 0.158 | 0.159 | 0.069 | 0.071 | 0.031 | 0.065 | 0.024 | 0.039 | |
| | | | | $Cyp46a1^{-/-}$ (n=3) | Mean | LLD | LLD | LLD | LLD | LLD | LLD | LLD | LLD | LLD | 5 |
| | | | | | SD | | | | | | | | | | |
| LMST01010164 | Cholest-5-ene-3β,24R-diol (24R-Hydroxycholesterol) | $C^{5}$-3β,24R-diol | 24R-HC | WT (n=3) | Mean | NQ | NQ | NQ | NQ | NQ | NQ | NQ | NQ | NQ | 6 |
| | | | | | SD | | | | | | | | | | |
| | | | | $Cyp46a1^{-/-}$ (n=3) | Mean | 0.004 | 0.005 | 0.012 | 0.005 | 0.009 | 0.013 | 0.009 | 0.018 | 0.023 | |
| | | | | | SD | 0.002 | 0.001 | 0.004 | 0.001 | 0.001 | 0.003 | 0.002 | 0.001 | 0.001 | |
| LMST01010018 | Cholest-5-ene-3β,25-diol (25-Hydroxycholesterol) | $C^{5}$-3β,25-diol | 25-HC | WT (n=3) | Mean | ND | ND | ND | ND | ND | ND | ND | ND | ND | 7 |
| | | | | | SD | | | | | | | | | | |
| | | | | $Cyp46a1^{-/-}$ (n=4) | Mean | 0.005 | 0.008 | 0.008 | 0.008 | 0.010 | 0.007 | 0.008 | 0.015 | 0.023 | |
| | | | | | SD | 0.002 | 0.001 | 0.001 | 0.002 | 0.003 | 0.004 | 0.003 | 0.017 | 0.034 | |
| LMST01010049 | 3β-Hydroxycholest-5-en-7-one (7-Oxocholesterol, 7-Ketocholesterol) | $C^{5}$-3β-ol-7-one | 7-OC | WT (n=3) | Mean | 0.002 | 0.002 | 0.002 | 0.002 | 0.003 | 0.003 | 0.004 | 0.002 | 0.002 | |
| | | | | | SD | 0.002 | 0.002 | 0.002 | 0.003 | 0.004 | 0.004 | 0.006 | 0.003 | 0.003 | |
| | | | | $Cyp46a1^{-/-}$ (n=3) | Mean | 0.002 | 0.003 | 0.002 | 0.002 | 0.003 | 0.002 | 0.003 | 0.002 | 0.002 | |
| | | | | | SD | 0.000 | 0.001 | 0.001 | 0.001 | 0.001 | 0.001 | 0.002 | 0.001 | 0.002 | |
| LMST04030168 | Cholest-5-ene-3β,7α,24-triol (7α,24-Dihydroxycholesterol) | $C^{5}$-3β,7α,24-triol | 7α,(24/25/26)-diHC | WT (n=0) | Mean | NA | NA | NA | NA | NA | NA | NA | NA | NA | 2 |
| LMST04030166 | Cholest-5-ene-3β,7α,25-triol (7α,25-Dihydroxycholesterol) | $C^{5}$-3β,7α,25-triol | | | SD | | | | | | | | | | |
| No ID | Cholest-5-ene-3β,7α,(25R)26-triol (7α,(25R)26-Dihydroxycholesterol,7α,27-Dihydroxycholesterol) | $C^{5}$-3β,7α,26-triol | | $Cyp46a1^{-/-}$ (n=3) | Mean | 0.003 | 0.002 | 0.002 | 0.002 | 0.004 | 0.003 | 0.004 | 0.002 | 0.001 | |
| | | | | | SD | 0.002 | 0.001 | 0.001 | 0.001 | 0.002 | 0.003 | 0.004 | 0.001 | 0.001 | |
| LMST04030170 | 7α,24-Dihydroxycholest-4-en-3-one | $C^{4}$-7α,24-diol-3-one | 7α,(24/25/26)-diHCO | WT (n=3) | Mean | 0.001 | 0.000 | 0.001 | 0.006 | 0.000 | 0.000 | 0.000 | 0.000 | 0.000 | |
| LMST04030107 | 7α,25-Dihydroxycholest-4-en-3-one | $C^{4}$-7α,25-diol-3-one | | | SD | 0.001 | 0.000 | 0.000 | 0.011 | 0.000 | 0.000 | 0.000 | 0.000 | 0.000 | |
| No ID | 7α,(25R)26-Dihydroxycholest-4-en-3-one (7α,27-Dihydroxycholest-4-en-3-one) | $C^{4}$-7α,26-diol-3-one | | $Cyp46a1^{-/-}$ (n=3) | Mean | 0.000 | 0.000 | 0.000 | 0.000 | 0.000 | 0.000 | 0.000 | 0.000 | 0.000 | |
| | | | | | SD | 0.000 | 0.000 | 0.001 | 0.000 | 0.000 | 0.000 | 0.000 | 0.000 | 0.001 | |
| No ID | 3β,7α-Dihydroxycholest-5-en-(25R)26-oic acid + 7α-Hydroxy-3-oxocholest-4-en-(25R)26-oic acid | $CA^{5}$-3β,7α-diol + $CA^{4}$-7α-ol-3-one | 3β,7α-diHCA + 7αH,3O-CA | WT (n=3) | Mean | 0.007 | 0.002 | 0.003 | 0.002 | 0.003 | 0.003 | 0.003 | 0.018 | 0.006 | |
| | | | | | SD | 0.010 | 0.002 | 0.002 | 0.002 | 0.001 | 0.002 | 0.002 | 0.009 | 0.002 | |
| No ID | | | | $Cyp46a1^{-/-}$ (n=4) | Mean | 0.002 | 0.002 | 0.002 | 0.001 | 0.003 | 0.005 | 0.004 | 0.013 | 0.005 | |
| | | | | | SD | 0.002 | 0.001 | 0.001 | 0.001 | 0.001 | 0.004 | 0.004 | 0.014 | 0.005 | |
| No ID | 7α-Hydroxy-3-oxocholest-4-en-(25R)26-oic acid | $CA^{4}$-7α-ol-3-one | 7αH,3O-CA | WT (n=3) | Mean | 0.001 | 0.001 | 0.001 | 0.002 | 0.002 | 0.001 | 0.001 | 0.003 | 0.002 | |
| | | | | | SD | 0.001 | 0.002 | 0.001 | 0.003 | 0.002 | 0.001 | 0.001 | 0.001 | 0.002 | |
| | | | | $Cyp46a1^{-/-}$ (n=3) | Mean | 0.000 | 0.000 | 0.001 | 0.001 | 0.001 | 0.001 | 0.001 | 0.001 | 0.001 | |
| | | | | | SD | 0.000 | 0.000 | 0.001 | 0.000 | 0.000 | 0.001 | 0.000 | 0.001 | 0.001 | |
| LMST01010001 | Cholest-5-en-3β-ol (Cholesterol) | $C^{5}$-3β-ol | Cholesterol | WT (n=3) | Mean | 305.1 | 366.8 | 446.8 | 252.9 | 356.7 | 575.5 | 421.3 | 263.3 | 509.2 | |
| | | | | | SD | 53.8 | 4.7 | 51.7 | 27.3 | 74.8 | 122.0 | 61.6 | 71.3 | 83.0 | |
| LMST01010016 | Cholest-5,24-dien-3β-ol (Desmosterol) | $C^{5,24}$-3β-ol | Desmosterol | WT (n=3) | Mean | 1.343 | 2.684 | 1.893 | 1.792 | 1.456 | 4.212 | 1.960 | 1.025 | 2.518 | |
| | | | | | SD | 0.334 | 0.027 | 0.097 | 0.085 | 0.072 | 1.546 | 0.613 | 0.356 | 0.162 | |
| LMST01010242 | Cholest-5,8(9)-dien-3β-ol (8(9)-Dehydrocholesterol) | $C^{5,8(9)}$-3β-ol | 8-DHC | WT (n=3) | Mean | 0.427 | 0.499 | 0.513 | 0.382 | 0.335 | 0.851 | 0.446 | 0.344 | 0.663 | |
| | | | | | SD | 0.089 | 0.071 | 0.117 | 0.069 | 0.083 | 0.427 | 0.050 | 0.120 | 0.023 | |
| LMST01010015 | Cholest-4-en-3-one | $C^{4}$-3-one | | WT (n=3) | Mean | 0.193 | 0.188 | 0.208 | 0.318 | 0.256 | 0.165 | 0.189 | 0.214 | 0.146 | |
| | | | | | SD | 0.228 | 0.190 | 0.175 | 0.304 | 0.276 | 0.142 | 0.199 | 0.144 | 0.167 | |
| | | | | $Cyp46a1^{-/-}$ (n=3) | Mean | 0.160 | 0.191 | 0.156 | 0.237 | 0.233 | 0.161 | 0.212 | 0.229 | 0.197 | |
| | | | | | SD | 0.026 | 0.034 | 0.032 | 0.028 | 0.094 | 0.027 | 0.104 | 0.019 | 0.062 | |

**Note.**

Abbreviations. LLD, Lower limit of detection; n, number of mice analysed; NA, Not analysed; ND, Not detectedd; No ID, no Lipid Maps ID; NQ, Detected but not quantified.

1. 24S,25-EC isomerises to 24-OC during the EADSA process.
2. Not analysed.
3. No authentic standard, identification based on accurate mass, $MS^{n}$ and retention time. Other possible identification (25S)26-hydroxycholesterol.
4. Approximate quantification by MRM and $MS^{3}$.
5. At or below lower limit of detection, $\leq$0.001 ng/mm$^{2}$.
6. Detected, not quantified.
7. Not detected.
